# Efficient Design of Peptide-Binding Polymers Using Active Learning Approaches

**DOI:** 10.1101/2021.12.17.473241

**Authors:** A. Rakhimbekova, A. Lopukov, N. Klyachko, A. Kabanov, T.I. Madzhidov, A. Tropsha

## Abstract

Active learning (AL) has become a subject of active recent research both in industry and academia as an efficient approach for rapid design and discovery of novel chemicals, materials, and polymers. The key advantages of this approach relate to its ability to (i) employ relatively small datasets for model development, (ii) iterate between model development and model assessment using small external datasets that can be either generated in focused experimental studies or formed from subsets of the initial training data, and (iii) progressively evolve models toward increasingly more reliable predictions and the identification of novel chemicals with the desired properties. Herein, we first compared various AL protocols for their effectiveness in finding biologically active molecules using synthetic datasets. We have investigated the dependency of AL performance on the size of the initial training set, the relative complexity of the task, and the choice of the initial training dataset. We found that AL techniques as applied to regression modeling offer no benefits over random search, while AL used for classification tasks performs better than models built for randomly selected training sets but still quite far from perfect. Using the best performing AL protocol, we have assessed the applicability of AL for the discovery of polymeric micelle formulations for poorly soluble drugs. Finally, the best performing AL approach was employed to discover and experimentally validate novel binding polymers for a case study of asialoglycoprotein receptor (ASGPR).

## 1. Introduction

Machine learning (ML) methods have been successfully applied in many areas of chemical research [1–5]. Classical ML models are trained on labeled datasets and then applied for virtual screening of chemical libraries to find hits with desired properties. However, in many areas of chemistry it is quite challenging to find a labeled dataset of sufficiently large size to enable rigorous building and external validations of property prediction ML models, as experiments are too expensive and slow to collect enough data. These problems have given rise to the field of active learning (AL) algorithms [6].

Active learning is an iterative procedure in which a machine learning model proposes candidates for testing to the user, and the user then returns labeled candidates, which are then used to update the model (Figure 1.). Usually, the main task of AL is to maximize the predictive performance of models using minimal required training data [6,7] and small datasets as user feedback. As applied to molecular and materials design, AL was shown to accelerate the experimental discovery of promising candidates by rapidly optimizing the molecular properties of interest [8–11].

**Figure 1.**
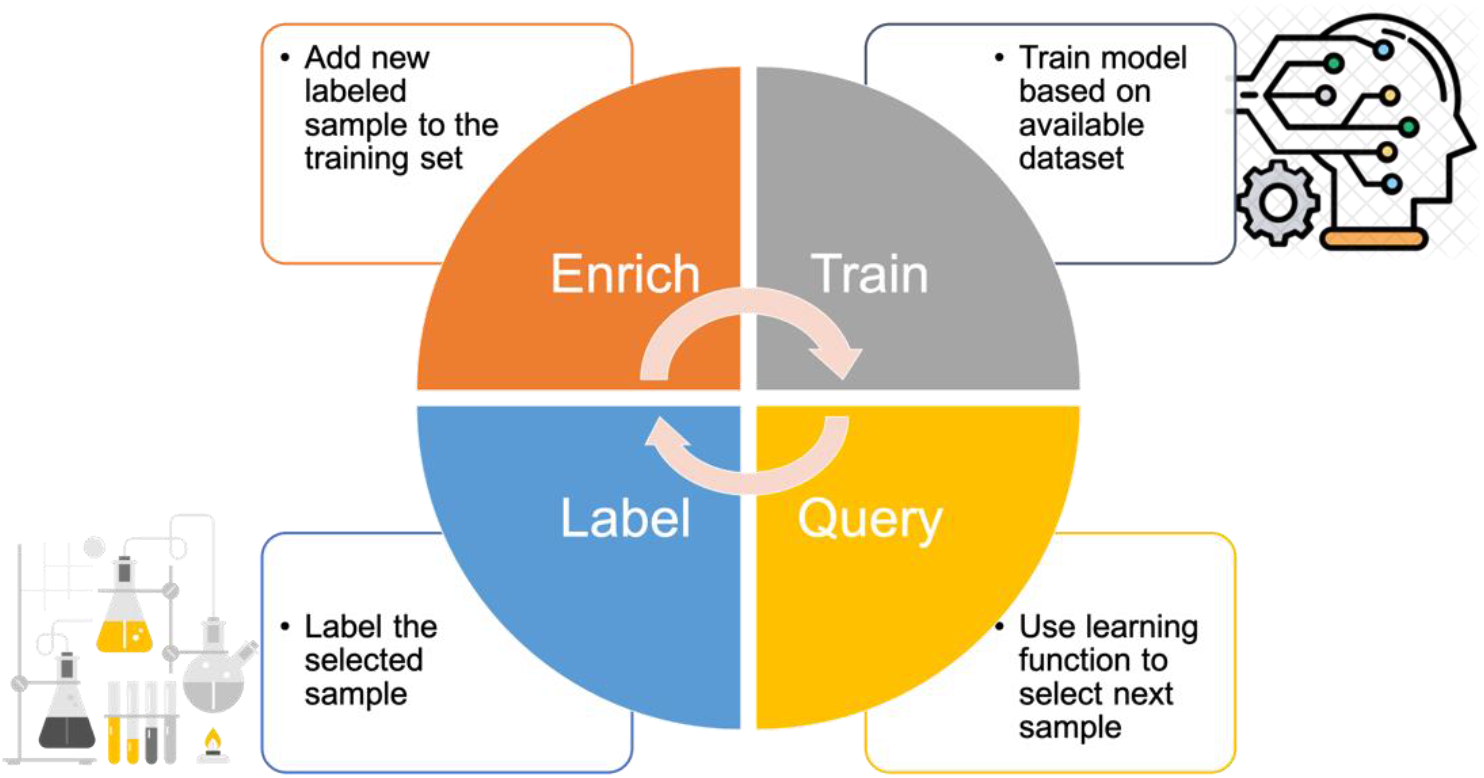
Active learning concept

New candidates for testing in the AL framework are either generated *de novo* if computational predictions are iterated with experimental testing or selected from a pool of accessible data [6]. Candidates are typically prioritized for testing based on some expected benefit for improving the predictive performance of the models. In applications of AL for chemical property optimization, in most cases the candidates are selected from unexplored regions of chemistry space where model uncertainty is the largest, which is referred to as “*exploration*” [7]. Alternatively, AL can be directed towards candidates with desired properties by maximizing the task-specific utility of a particular action/measurement [8–12]; this approach is referred to as “*exploitation*”.

There has been a growing interest both in basic and applied research in exploring AL as a means of efficient search, design, and discovery of chemicals, materials, and polymers. For instance, there have been several recent efforts on the use of AL strategies to predict reaction yields [8], discover organic semiconductors [13], search for polymer dielectrics with a large band gap [14], design novel ^19^F magnetic resonance imaging (MRI) agents [15], develop ML models that predict quantum chemical properties [16–19], and facilitate drug discovery [10,11,20] (reviewed recently by Reker et al. [21]).

In all aforementioned examples, initial datasets were usually quite large and only several studies were reported where AL started from extremely small datasets [9,19]. Kim et al. [9] evaluated the effectiveness of three AL strategies (exploitation, exploration, and the balanced exploitation and exploration method) compared to a random selection (when the training dataset is selected randomly) approach for the detection of polymers with high glass transition temperatures (Tg > 450K). The initial surrogate model was built for five randomly selected polymers using Gaussian process regression (GPR). They found that the balanced exploration-exploitation approach had the highest accuracy and efficiency to identify polymers with desired properties. Loeffler et al. [19] attempted to minimize the size of the training set to build neural network models for predicting energy of water clusters. They introduced AL strategy that starts with minimal training data and is continuously updated via a nested ensemble Monte Carlo scheme. The authors generated training data on the fly by selectively adding configurations from failed regions. They showed that this strategy can be applied to develop accurate force-fields for molecular simulations using sparse training data sets.

Herein, we report on an AL methodology for efficient and quick search for candidate compounds with the desired properties using minimal data and limited experimental testing. Following the initial benchmarking studies of different AL protocols, we employ the best-performing AL approach to the discovery of synthetic polymers that can selectively bind specific peptides or small molecules. Such polymers can be used as synthetic scavengers of xenobiotics [22,23], as micelles for targeted delivery of peptides or small molecules [24] or to enhance solubility of poorly soluble drugs [25], to name a few. Due to lack and sparsity of data, classic QSAR modeling cannot be applied in this case. Thus, this study focuses on the design of polymeric systems in the common case of small training datasets.

We chose the asialoglycoprotein receptor (ASGPR) as a target for molecular recognition screening. ASGPR is involved in the endocytosis of the desialylated glycoproteins. The receptor is located predominantly on the surface of hepatocytes and is also found on urethral epithelial cells [26] and human sperm cells [27]. It is actively expressed by hepatocellular carcinoma (HCC) cells [28]. It belongs to the C-type lectin family, which also includes mannose macrophage receptor, Dendritic Cell-Specific Intercellular adhesion molecule-3-Grabbing Non-integrin, subset of selectins (E-selectin, P-selectin, L-selectin), and a number of others [29,30]. ASGPR takes part in the pathogenesis of the Marburg hemorrhagic fever [31], viral hepatitis [32], and gonorrhea [26]. It is also involved in the development of colon adenocarcinoma metastasis [33], as well as in the metastasis of other cancers by activation of EGFR – ERK with upregulation of MMP-9 [34]. ASGPR was targeted for the delivery of anticancer therapy to HCC cells [35–38] and for delivery of the copper chelation agents as a therapy for Wilson disease [39]. ASGPR was used for identification of the ligand epitope and design of polyplex-based gene carrier [40]. Other examples of ASGPR-based targeted delivery include the treatment of acute hepatic porphyria [41], viral hepatitis [42,43]. It was also involved in the delivery of atorvastatin conjugated with the targeting moiety into hepatocytes for up-regulation of cholesterol metabolism in hypercholesterolemia patients [44], providing reduction of systemic distribution-related side effects, such as myopathy and muscle pain [45]. ASGPR was used as the model target for the development of neuronal gene delivery platform. Even though the ASGPR is not expressed by the primary sensory neurons, the detection of several galectin receptors with similar ligand profile was observed.

Thus, due to its participation in various biological processes, ASGPR represents an attractive model object for creating a molecular recognition system. Although there are recognition systems based on small molecules, the macromolecule-based recognition system can exhibit greater specificity and stability. Macromolecules and polymers can interact with the cell membrane and the receptor in the fashion different from that of low molecular weight compounds. The biodistribution and pharmacokinetic profile are expected to be different as well, offering the opportunity to overcome the problems typical for small molecule-based drugs. A novel polymer-based targeting moiety could be conjugated with the active pharmaceutical ingredient molecule for directed treatment of various liver diseases. Meanwhile, the water-soluble polymer-based recognition system can prevent the ASGPR-targeted interaction of various pathogens, such as viruses (Marburg virus, Hepatitis A and B viruses) or bacteria (*Neisseria gonorrhoeae*).

This paper is organized as follows. First, we conduct a series of synthetic experiments to study and compare various AL strategies using a small initial training set (5 to 50 data points) with known values of biological activities (pIC50, pKi) of chemical compounds. In doing so, the goal is to accelerate the discovery of the desired compounds, which is the ultimate objective of the experimental research, rather than improve the overall model performance. Second, we investigate how the performance of each AL strategy may vary depending on (i) the size of the initial training set, (ii) different approaches to selecting the initial data, and (iii) the relative complexity of the task at hand. Finally, we evaluate the proposed AL methodology for finding polymers that can bind organic molecules based on the literature data [46] and using real experimental setup as well as polymers that can bind ASGPR.

## 2. Materials and methods

### 2.1. Datasets and descriptors

#### 2.1.1. ChEMBL datasets

The comparison and selection of optimal AL approaches was carried out using datasets with known values of biological activities (pKi) of chemical compounds tested in biological assays for different target proteins; all the data was downloaded from the ChEMBL database (CHEMBL205, CHEMBL244, CHEMBL217) [47]. A histogram of the distribution of pKi values in the selected datasets is shown in Figure 2.

**Figure 2.**
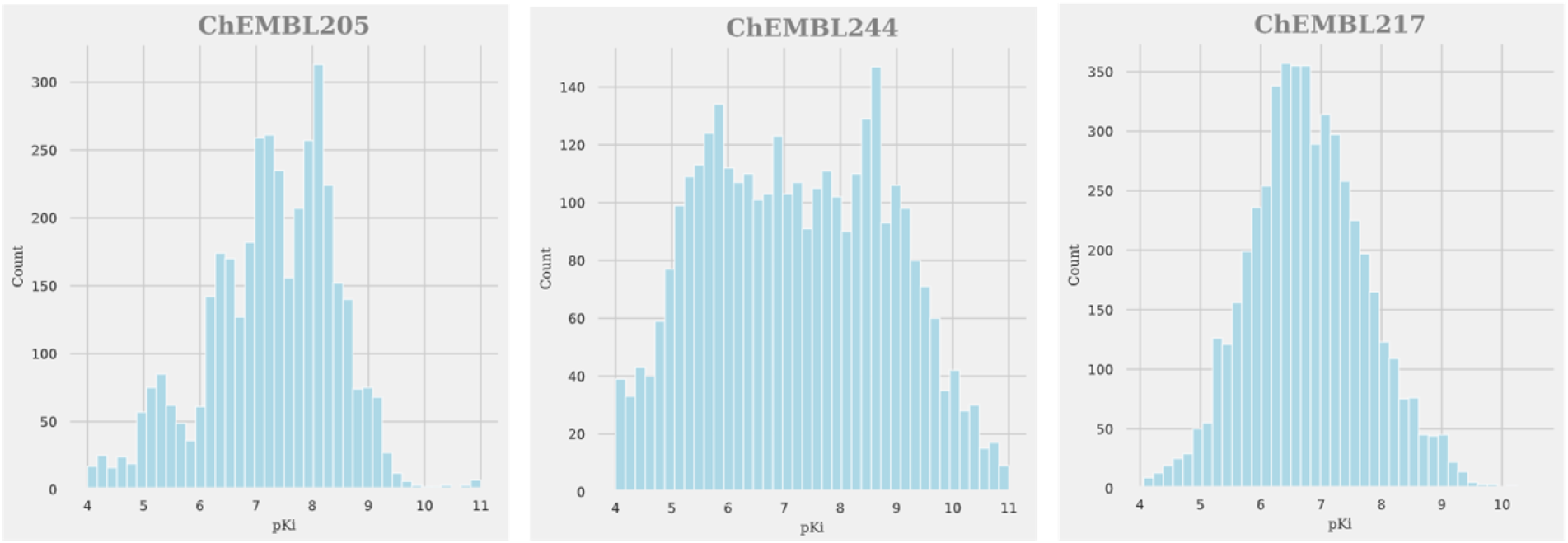
Distribution of the pKi values for all the datasets considered in this study

Molecules were randomly divided into training (80%) and external test (20%) sets (Table 1). Regression models were built using pKi values whereas to build binary classification models, we assumed that molecules with pKi greater than 7 were active, otherwise inactive (the distribution of active and inactive molecules is presented in Table 1).

**Table 1.** Sizes of training and external test sets for three datasets. Number of active/inactive compounds is given in brackets

| Dataset | Size of dataset | Size of training set | Size of external test set |
| --- | --- | --- | --- |
| CHEMBL205 | 3808 (2463/1345) | 3047 (1977/1070) | 761 (486/275) |
| CHEMBL217 | 5011 (2016/2995) | 4009 (1607/2402) | 1002 (409/593) |
| CHEMBL244 | 3305 (1771/1534) | 2644 (1403/1241) | 661 (368/293) |

##### Descriptors

Morgan fingerprints with a size of 1024 bits and a radius of 2 generated using the RDKit package [48] were used as descriptors for modeling.

#### 2.1.2. Polymer datasets

We used literature data [46] to assess the application of AL strategies for the discovery of polymers with specified binding affinities for organic molecules. We have employed published data [46] on the loading capacity (LC) of five hydrophobic drugs (Curcumin (CUR), Paclitaxel (PTX), antiretroviral efavirenz (EFV), Dexamethasone (DEX), anti-oxidative tanshinone IIA (T2A)) for micelles formed by 18 different amphiphilic polymers. The loading capacity values were measured at different time intervals: 0, 5, 10, and 30 minutes after adding the drug to the polymer solution. LC for PTX, EFV and CUR was measured at all time points; and LC for DEX and T2A were measured only at 0 minutes. Thus, our collection included 252 data points for drug-polymer systems. For this dataset, the high value of loading capacity (LC > 33%) was observed for 77 drug-polymer systems. Such points were considered as having class label “active”, or “1”, and the remaining systems were assigned to class “0”. Models were developed for eight different datasets, and every dataset contained LCs data for a given drug, measured at a given time interval.

Only the solubilization data for curcumin (CUR) and paclitaxel (PTX) contained instances of active and inactive classes and therefore these datasets were selected for model development (other drugs were insoluble in the presence of any polymeric micelle; see Table S1, Supporting Materials). The stability of polymer-drug composition with CUR and PTX drugs after certain time (0, 5, 10, and 30 minutes) were used for testing AL approaches. Thus, we had 8 datasets with 18 datapoints in each: a dataset included data for one drug (CUR or PTX), solubilized with 18 different polymeric micelles for one of the four time intervals. Then, an individual virtual experiment was conducted to design an optimally solubilizing polymer.

For experimental validation, AL workflow was used for the design of polymeric recognition system for ASGPR. The design of this system aimed to find block copolymers of PEG and polyamine acids (polylysine - PLKC, polyglutamate - PLE, polyaspartate - PLD) that bind to the model protein ASGPR. PEG block can have molecular weight of 5k or 1k; PLKS, PLD or PLE polymers can have 10, 30, 50, 100 repeated blocks. The α-end of the polyethylene glycol could have a CH_3_O group or an azide group. 13 different types of polymers were synthesized in the project (see Results section).

##### Descriptors

To describe the chemical structure of small molecules, polymers, and drug-polymer systems, we used modified simplex descriptors described in [25]. The descriptors for drug-polymer systems included:

i. traditional SiRMS descriptors (multiplets containing 4 atoms) of a pseudo small molecule representing a polymer. Pseudo small molecule is a block of polymer repeated only once with corresponding terminal blocks of the polymer. In case of block-copolymer both blocks are repeated once;
ii. “mixture” simplex descriptors of a drug-polymer system. In every simplex at least one of four atom comes from drug molecule and at least one comes from a polymer building block;
iii. composition descriptors, which includes the number of particular monomer repetitions in a polymer structure;
iv. time interval for the loading capacity measurement (0, 5, 10, or 30 minutes). Used only for the LC dataset.

In case of highly correlated (r ≥ 0.9) descriptors, one of them was chosen arbitrarily and removed; similarly, we removed low variance descriptors to reduce the dimensionality of the chemical space without losing important information. A total of 446 descriptors were obtained.

For the design of ASGPR polymeric recognition system descriptors of the target molecule were not added. Thus, only SiRMS descriptors [25] of the pseudo-small polymer molecule were taken to describe the polymer structure as described above. Low variance and highly correlated descriptors were removed. As a result, 21 SiRMS descriptors were obtained, which was augmented by 2 descriptors containing information about polymer composition (MW of PEG block and number of polyaminoacid block).

SiRMS descriptor calculation was performed using Python implementation accessible in GitHub: https://github.com/DrrDom/sirms. Descriptor selection was accomplished using built-in tools from scikit-learn library [49,50].

### 2.2. Modelling

Random Forest Regression (denoted as RFR), Classification (denoted as RFClf), and Gaussian Process Regression (denoted as GPR) approaches were employed for building quantitative structure-property relationships (QRPR) models. Implementations of the RFR, RFClf and GPR methods were taken from the scikit-learn library [50]. Random Forest is an ensemble of multiple decision trees trained independently on a random subset of training data. This method has a small number of hyperparameters and is insensitive to the presence of many descriptors. The predicted classification values are defined by the majority voting for one of the classes. The predicted regression values are defined by the mean prediction of the individual trees. As regression model’s prediction confidence, required for some AL approaches, we used variance of tree predictions in Random Forest. We slightly modified the original RFR code to extract variance together with prediction itself. The number of trees in RFR and RFClf was 500, the number of features selected upon tree branching (max_features option) was set to log2 of features number. Such settings have shown best performance in our tests. Other hyperparameters of RFR and RFClf were set to default values.

GPR assumes that the joint distribution of a real-valued property of chemical objects and their descriptors is multivariate normal (Gaussian) with the elements of its covariance matrix computed using special covariance functions (kernels). GPR model produces a posterior conditional distribution (so-called prediction density) of the property given the vector of descriptors for every chemical object. The prediction density has a normal (Gaussian) distribution with the mean corresponding to the predicted value of the property and the variance corresponding to prediction confidence [51]. Based on our tests, for GPR models hyperparameters of noise level, alpha was set to 0.1, and RBF kernel’s gamma value was 10. Other hyperparameters of GPR were set by default.

### 2.3. Testing different AL strategies using ChEMBL datasets

For testing different AL approaches, we used ChEMBL datasets described above. Our objective was to compare these strategies for their relative ability to find as many candidate compounds that possess desired property as possible.

#### Training and test datasets

The dataset applied for training surrogate model for the first time is called the *initial set*; it is usually quite small (5, 10, 20, 50 objects). After an AL cycle, the initial dataset is enriched by the fixed number of new data and the resulting dataset is called *current set*. As the AL cycles are executed progressively, the size of the current dataset gradually increases.

As the validation set for each model, we use the data that was left after exclusion of the current dataset data points from the training sets. All model quality metrics are measured on the external test set, which is selected in advance and by no means used in modeling (Figure 3).

**Figure 3.**
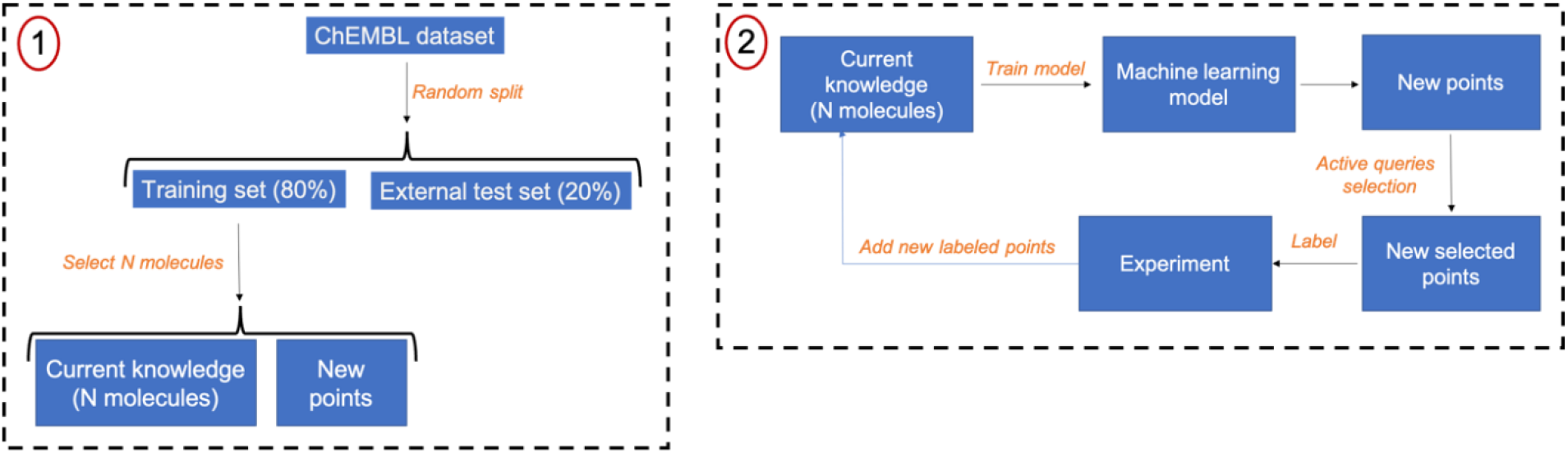
Illustration of AL method. Left - dataset division into training and external test sets. Right - overview of the AL algorithm. First, the AL model is trained based on current knowledge. Using the surrogate model, predictions, and associated uncertainties are obtained for new points. New points are selected based on respective AL strategies (see text). After testing new selected points, the results are used to update current knowledge, and iterations are repeated until the desired goal is achieved

#### Selection of the initial dataset

The active learning cycle starts from training the surrogate model on the initial dataset of N objects (N = 5, 10, 20, 50); see Figure 3. The initial dataset is comprised of data points that have to be experimentally measured prior to AL modeling. Two strategies for selecting the initial dataset were tested:

1. random selection (RANDOM) – N objects are randomly selected from a set of possible candidates;
2. selection of the most diverse objects (DIVERSE) - the entire sample is clustered by KMeans algorithm implemented in the scikit-learn library [50]. One random object is selected from each cluster to form the initial dataset. Thus, the number of clusters was set equal to the desired size of the initial set.

#### Active learning strategies

Initially, descriptors of selected objects for testing as well as value of their property are retrieved. Then we train the surrogate model on the *initial* set using the machine learning model that relates descriptors of the molecules and their pKi. After building the surrogate model, pKi values (along with associated uncertainties) or class probabilities are predicted for the remaining candidate molecules from the validation set. Based on these predictions, K (hereafter, K=1, 2 or 5) molecules are selected for further experimental testing and added to form the *current* set. The following selection strategies were tested:

- *Strategy 1. Exploitation* (Y-MAX) – Top K molecules with the highest predicted values are selected as new objects.
- *Strategy 2. Exploration* (Y-VAR) – objects with higher prediction uncertainties are selected. For regression, candidates with the maximum variance of predictions returned by RFR are selected. For RFClf, we calculated the variance of the predictions from the Bernoulli distribution equation as var=p(1-p), where p is probability that the molecule is active. So, the closer the probability to 0.5 the larger is the variance. The *exploration* approach is efficient in making the model more robust, but the added object may not be active [6,7].
- *Strategy 3. Hybrid selection* (Y-MAX(M)-Y-VAR(L)) – the best M = K-L candidates are selected according to the Y-MAX strategy, while the best L candidates are selected according to the Y-VAR strategy. This approach exhibits balance between exploration and exploitation. In the work, Y-MAX(2)-Y-VAR(3) or Y-MAX(1)-Y-VAR(1) is used.
- *Strategy 4. Random selection* - in this strategy, we randomly select new candidates at every iteration and add them to the initial or current set. This strategy serves as negative control.
- *Strategy 4. Hypothetical perfect strategy –* this approach approximates optimal selection strategy. To find perfect candidate for adding, we look for the candidate, addition of which to current set improves the predictive performance of the surrogate model the most. It cannot be considered as AL strategy, since we need to know actual property value for every candidate to select the best one (in AL we don’t know candidate property in advance). This approach is used for benchmarking purpose only.

The AL cycle was repeated continuously until the size of the *current* set reached 150 objects (Figure 3). The entire process was repeated 10 times for each strategy, each time starting from a different initial dataset to eliminate the bias associated with the initial selection of N points.

#### Statistical analysis

Once the surrogate model is built, its utility for selecting new objects is assessed on the external test set. The following statistical metrics were used to assess different aspects of performance of AL strategies:

1. ***Model accuracy performance*** – predictive performance of models obtained on a current set. Coefficient of determination R^2^ was used to characterize the accuracy of the regression models and was calculated as shown in classic formulae (1). Balanced accuracy BA was used to assess classification model performance and was calculated as shown in formulae (2). The motivation of this metric is that often AL is used to build best model with least amount of data [6,7]; also model with high predictive ability can better select candidates with desired property values for further testing.

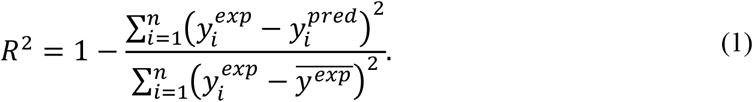

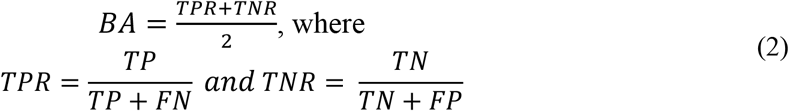
2. ***Candidate improvement performance***. This metric reflects the quality of the candidate selection process, i.e., how well the surrogate model selects highly active compounds from the external test set. The quality of selection was characterized using the Enrichment Faction criterion (EF_10_). For regression, it is equal to the fraction of 10% most active compounds in the list of the 10% most active compounds as predicted by model. For classification, EF_10_ was equal to the fraction of 10% most active compounds (selected according to pKi values) in the list of 10% molecules with the highest probability of positive class (includes only objects with probability greater than 0.5). Thus, if the model correctly retrieves most active compounds, EFis equal to the maximum value of 1.

### 2.4. Experimental settings in the ASPGR binding measurement

The experiment was conducted on a Biacore X100 machine (Biacore AB, Uppsala, Sweden) using a CM5 chip. ASGPR from rabbit liver was purchased from Generic Assays (GA Generic Assays GmbH, Berlin, Germany) and used for SPR assays. The ASGPR was immobilized according to the standard amine-coupling protocol provided by the manufacturer. Polymer ligands were dissolved in a running buffer (150 mM NaCl, 50 mM CaCl_2_, 50 mM Tris, pH 7.4) followed by serial dilution up to 5*10^−6^ M. Samples were injected at a flow rate of 20 μL/min at 25 °C for 30 s followed by 30 s dissociation. The regeneration of the sensor chip was obtained by injection of 20 μL of 20 mM EDTA. All solutions were filtered and deoxygenated. Data were analyzed using BIAevaluation 3.0 software. The K_D_ values were evaluated using the 1:1 Langmuir association model. Data are mean ± SD of n = 3 independent measurements.

## 3. Results and discussion

### 3.1. Active learning strategy for finding biologically active molecules in “virtual” experiments

To compare different AL strategies introduced above, we simulated AL workflow using “virtual” experiments, represented by the collection of datasets with known values of biological activities (pKi) of chemical compounds.

#### 3.1.1. Optimization of continuous property

First, the performance of AL using regression modelling for optimization of continuous properties was tested. Table 2 demonstrates the performances of RFR and GPR models trained on the entire training set. The performances of regression models are quite similar.

**Table 2.**
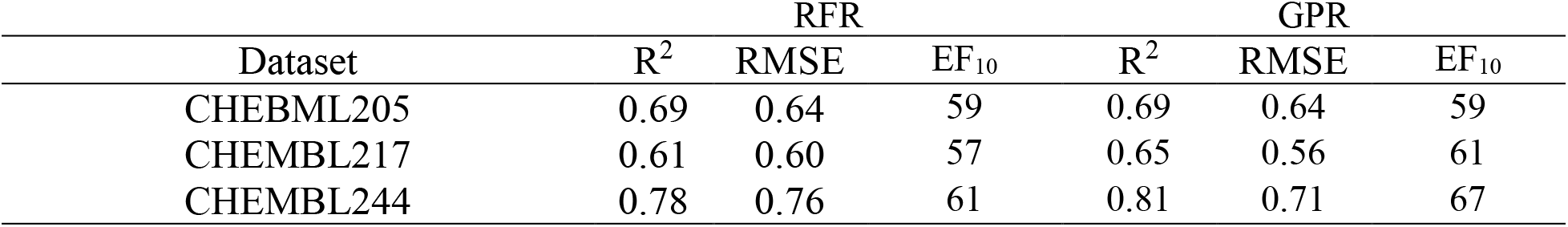
Coefficient of determination (R^2^), RMSE of predictions, and EF_10_ estimated on the external test set using RFR and GPR models trained on the entire training sets

We analyzed the influence of different strategies (Y-MAX, Y-VAR, and *Random selection*) on the performance of the AL algorithms for RFR and GPR models (see Supporting Materials, Figure S1-S2). In general, RFR and GPR machine learning approaches were characterized by quite similar change in the model accuracy (R^2^) and virtual screening (EF_10_) metrics as the dataset size was increased. Thus, only RFR is used below due to its simplicity and speed. The maximum value of the coefficient of determination achieved 0.38 for 150 iterations, the maximum values of EF_10_ were about 30-40% which is much lower than corresponding metrics for model trained on all data (shown in Table 2). None of machine learning approaches combined with Y-MAX, Y-VAR AL strategies was better than random selection within 150 iterations. Notably, Y-VAR strategy was almost always better than Y-MAX strategy.

In the experiments above, the AL cycle stopped when 150 compounds were selected, which can be insufficient for modeling; thus, we tested AL strategies by adding compounds iteratively until current set contains the whole training set (Figure 4). It was found that to achieve 70% value of the model’s performance built on the entire dataset, more than 1000-1500 points must have been added to current set. Moreover, the R^2^ of considered AL strategies was never better than *Random selection* (red line in Figure 4) showing no benefits of AL application in this case. Interestingly, despite quite poor R^2^ values, Y-MAX strategy shows EF_10_ curve above the line of Random selection in CHEMBL217 dataset [47]. Since our goal was to find the desired compounds with minimal experimental effort, Y-MAX AL strategy is a reasonable alternative to random search. In general, for datasets explored in this study, we saw no benefits of AL application in regression modeling. Possible reason for this observation can be related to the selection of noisy datasets with high random measurement error collected from different bioassays. Such a large fluctuations of predicted value could mimic true difference in activity values for compounds, especially if small datasets are used.

**Figure 4.**
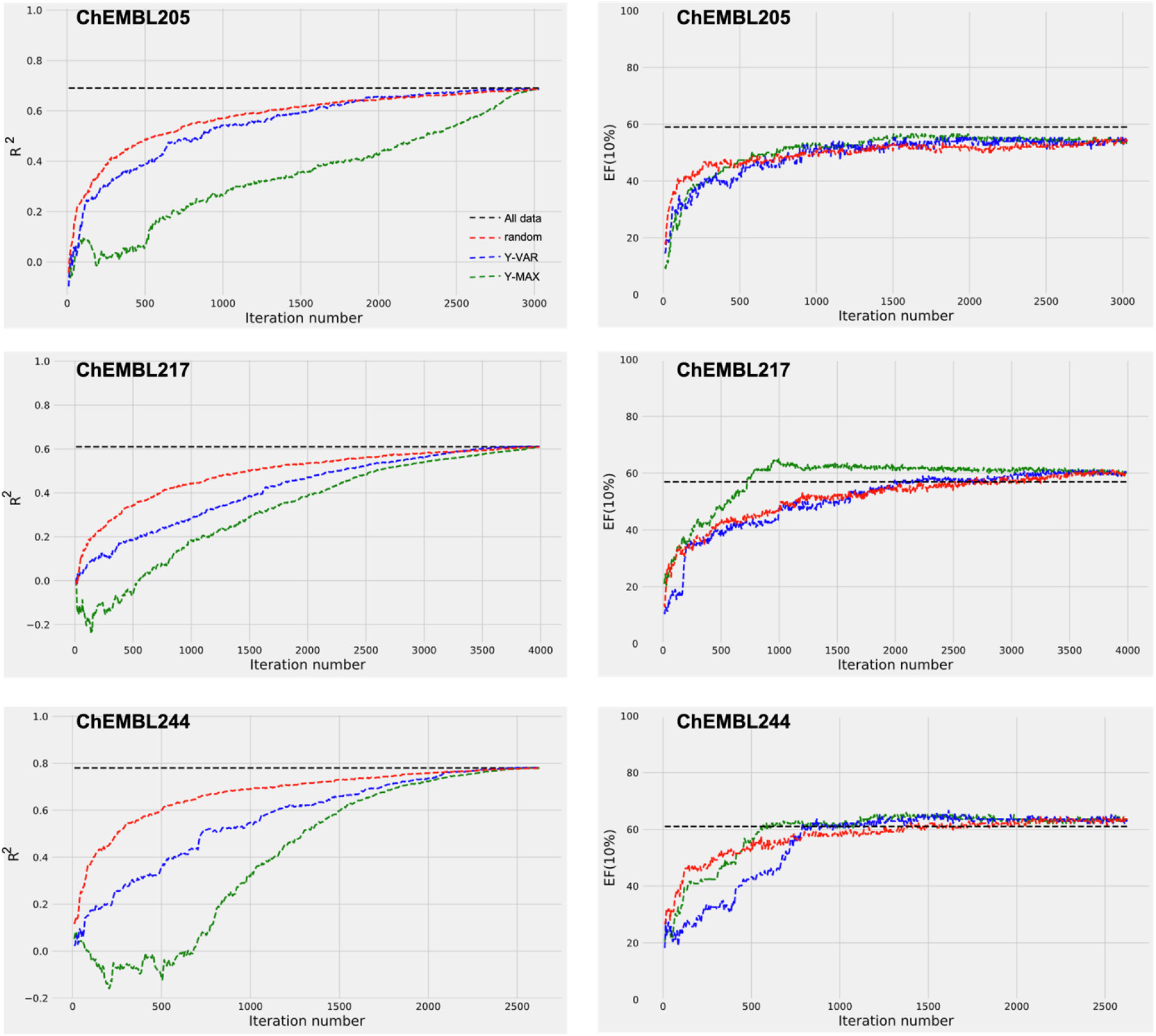
Influence of different AL strategies on the quality of RFR. The initial set contained 10 points; 5 objects were added at every AL cycle. Active learning strategies are Y-MAX (green line), Y-VAR (blue line), and random (red line), black dotted line represents performance of the model trained on the entire training data set. Lines correspond to median values over 10 repetitions

#### 3.1.2. Binary classification models

To reduce noise in pKi values, which could lead to weak model performance in AL iterations, classification modeling was selected for surrogate model building. Compounds with pKi greater than 7 were considered active; otherwise, they were labelled as inactive. Models built with conventional QSAR approach using complete training sets were characterized by reasonably high balanced accuracy (more than 0.8, Table 3).

**Table 3.**
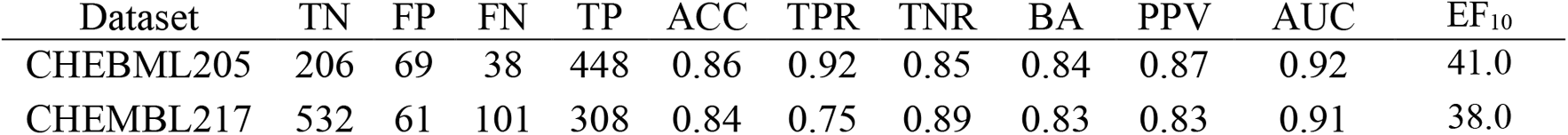
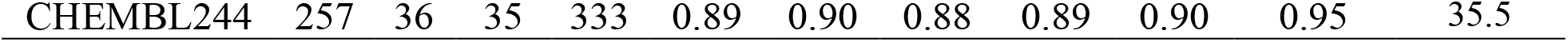
The predictive performance of the external test sets using RFClf models trained on all training sets

The influence of different strategies (Y-MAX, Y-VAR, and Random selection) on the performance of the AL algorithms for RFClf models is shown in Figure 5-Figure 7. While in the case of regression modeling the predictive performance of models was poor (R^2^ ∼ 0) when only small number of up to 150 datapoints was explored (cf. Figure 4), in the set used for classification model building Y-VAR performed better than regression models even if datasets were small (BA is significantly greater than 0.5, Figure 5). In contrast, Y-MAX strategy had mediocre BA on small datasets without trend to grow when up to 150 datapoints in the current set were explored (Figure 5). On the contrary, for the regression models, we have higher values of EF_10_ than for classification models even if all training data are used for modeling (black lines in Figure 4 and Figure 5). Thus, continuous scale of predicted values helps model to better select most active compounds. It is quite logical since classifier has only class labels without information on actual activity values.

**Figure 5.**
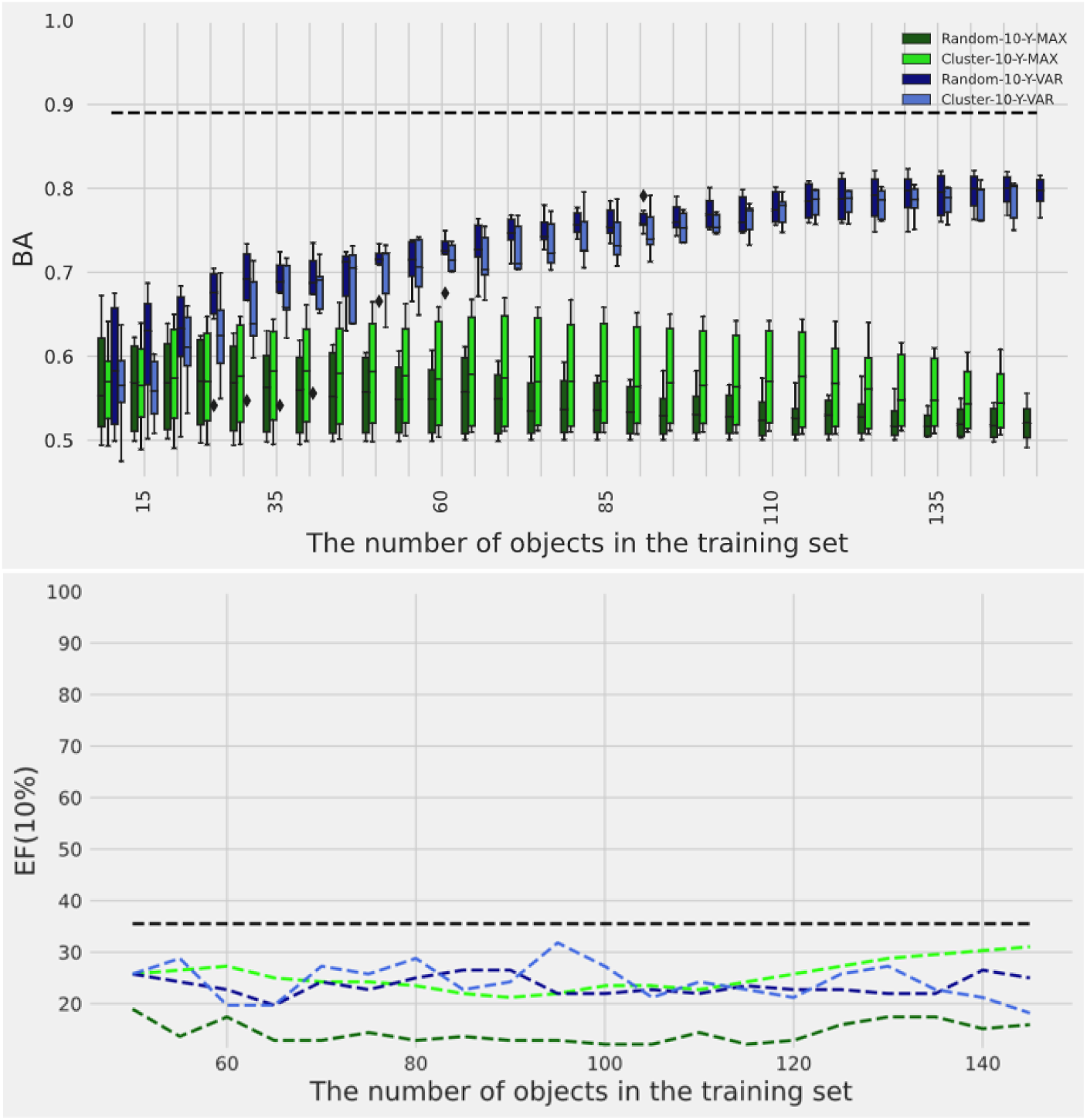
Influence of different selection protocols for the initial dataset on AL performance (for CHEMBL244 dataset). Initial set was selected randomly (Random, deep green or deep blue) or as diverse set using clusterization (Cluster, light green, or light blue). Y-MAX (deep and light green) and Y-VAR (deep and light blue) strategies were tested. The initial set contains 10 points, 5 objects were added at every AL cycle. Black dotted line represents performance of the model trained on entire training data set. Statistics corresponds to 10 repetitions of AL cycle; lines correspond to median values

We analyzed the effect of the initial data selection procedure on the performance of each AL strategy (Figure 5, the results for other sizes of the initial training set and for other datasets are given in Figure S3-S6 of the Supporting Materials). It turned out that the formation of the initial dataset from the most diverse objects did not have an advantage over random selection. This conclusion was valid for all datasets, different sizes of the initial training set, and different AL strategies (Y-MAX, Y-VAR). Thus, below we will demonstrate results for the simplest method, a random initial set selection. The size of initial dataset also influences model performance. In general, larger initial sets enable models with greater performance. BA values at different initial set size is shown in Figure 6 for CHEMBL244 dataset. For Y-MAX strategy it is better to start from a larger dataset, since as is shown above (Figure 4), Y-MAX selection leads to worse model than random selection of the same number of molecules. On the contrary, using Y-VAR strategy one may start with smaller number of objects in initial dataset iteratively adding objects to it, and the model performance would be similar (Figure 7).

**Figure 6.**
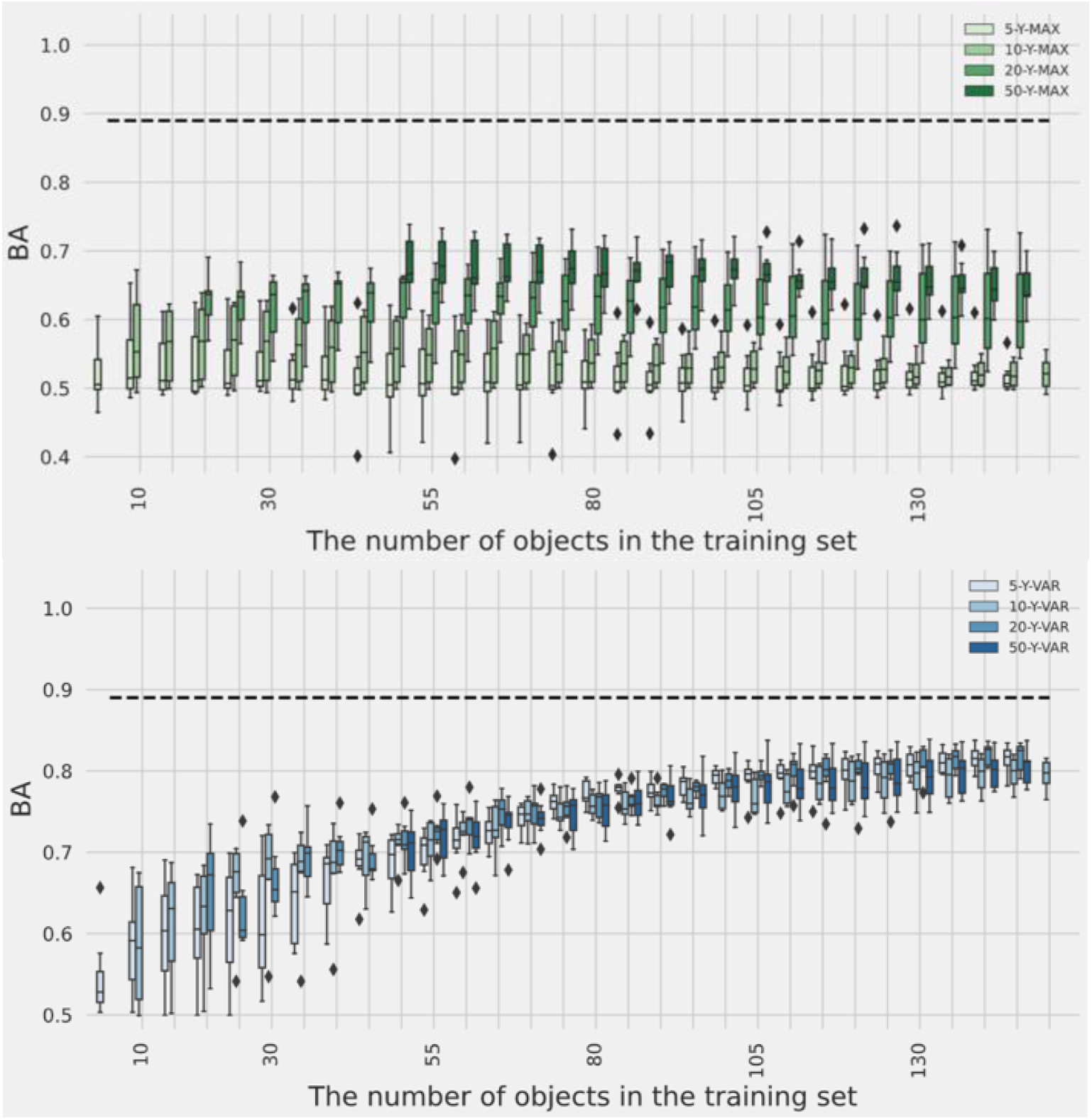
Influence of the initial dataset size (for CHEMBL244 dataset) on BA for the external test set. Y-MAX (top) and Y-VAR (bottom) strategies were tested. Five objects were added at every AL cycle. Black dotted line represents performance of the model trained on the entire training data set. Error bars are calculated by averaging the results of 10 repetitions of the AL selection cycle

The influence of different AL strategies: Y-MAX, Y-VAR, *Hybrid* Y-MAX-Y-VAR has been thoroughly investigated in comparison with *Random selection* and *Hypothetical perfect strategy* (Figure 7). Notice, that hypothetical perfect model has both best BA values (closest to model built on all data), and highest EF_10_ among others. It supports intuitive concept that the more performant models are more efficient in compounds selection. As in the case of regression models, the Y- MAX strategy very slowly improves the performance of the classification models. The models were biased towards a positive class showing high TPR on external dataset (high TPR in Figure S7 Supporting Materials, close to 1.0), but low precision value (PPV in Figure S7), which is close to positive class fraction in the dataset. Also, we observe that EF_10_ values for Y-MAX were the worst among all other strategies. Only when current dataset size achieves 500 molecules or more, EF_10_ increases and the approach becomes the best in active compound selection with small advantage over *Random selection* (Figure S7), despite model still staying biased towards positive class.

**Figure 7.**
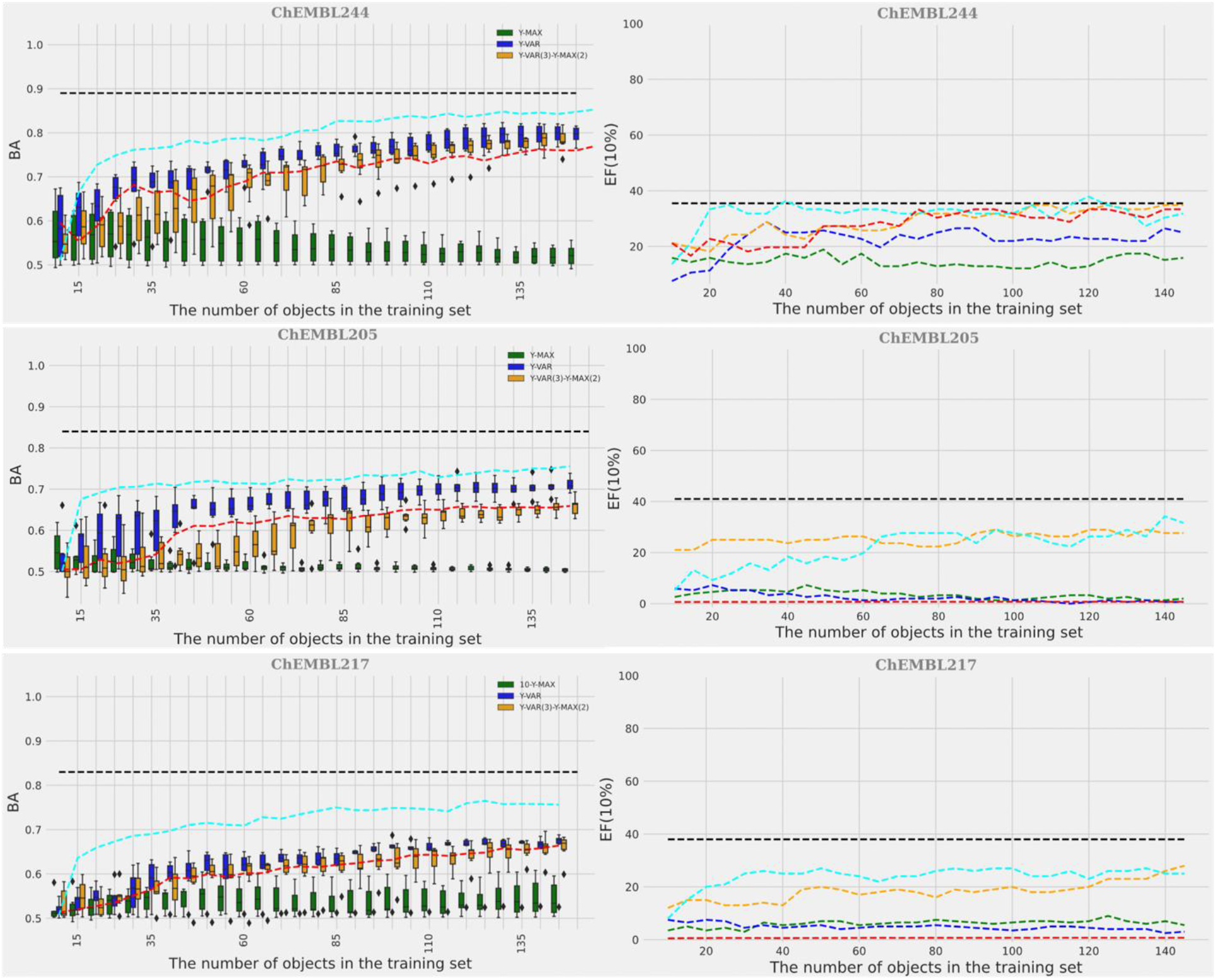
Performance of different AL strategies: Y-MAX (green box and line), Y-VAR (blue box and line), Y-MAX(2)-Y-VAR(3) (orange box and line), in comparison with random selection (red dotted line) and hypothetical ideal model (cyan dotted line). Lines correspond to median values over repetitions. The initial datasets consist of 10 points. Black dotted line represents performance of the model trained on entire training data set

As shown in Figure 7 and Figure 8, Y-VAR strategy and, to a lesser extent, *Hybrid* Y-MAX-Y-VAR strategy had an advantage over *Random selection* in improving the performance of the classification models (BA values on external test set). But when small current sets are selected (up to 150 objects, Figure 7), EF_10_ values for Y-VAR and *Hybrid* Y-MAX-Y-VAR strategies did not outperform models that relied on *Random selection*. Further selection by Y-VAR (shown in Figure S7) makes even worse and EF_10_ values were generally worse.

**Figure 8.**
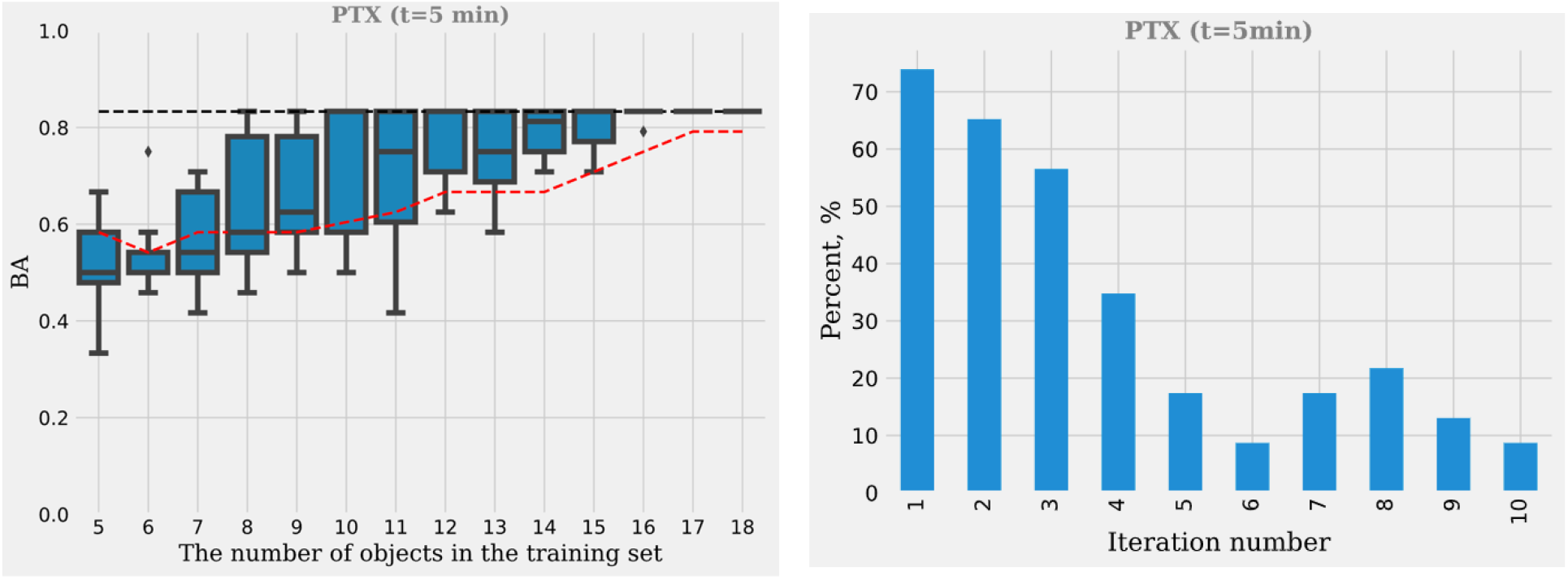
The performance of selection of the optimal polymer for PTX solubilization. Solubilizing ability was studied after 5 minutes of mixing. On the left - the predictive performance of the model built on current set on validation set, the BA value is given. On the right - the chance that selected polymer is active (solubilizing). Baseline BA value for the model built on all data assessed in leave-one-out validation is shown by black dotted line

*Hybrid selection* Y-MAX-Y-VAR in combination with classification models showed reasonable quality for ChEMBL datasets (Figure 7). It showed slightly lower BA values in comparison with Y-MAX, but still generally better than *Random selection*. At the same time, it has shown largest EF_10_ value, reflecting its utility for the selection of active compounds (Figure 7, top right). It is usually better than *Random selection* and approaches the accuracy that can be seen with *Hypothetical perfect strategy*; however, fluctuations are rather high. Thus, *Hybrid selection* Y-MAX-Y-VAR shows reasonable quality in both tests and adopts best features of Y-VAR strategy to enhance model quality and Y-MAX, which usually better selects active compounds.

To summarize, in our virtual experiment we have compared different AL approaches. Application of AL to continuous value properties required the selection of rather large datasets, but for binary property, AL strategies had shown good results even for small datasets. Neither machine learning method, nor the initial set selection strategy have influenced the accuracy of AL models. Conversely, the selection strategy affected the AL performance the most. Exploration (Y-VAR) and hybrid strategies could select compounds such that the model accuracy is higher than if compounds were selected randomly. Hybrid strategy also allows to more efficiently select compounds with the desired property, and thus we chose this approach for further application.

### 3.2. Testing active learning strategy in searching for polymers with good solubility

We tested AL strategy in a case study where we searched for polymers that form micelles used in drug delivery systems. The dataset with drug LC for different polymeric micelles was extracted from publication [46].

To form the initial training dataset, five random polymers were selected. The model was built using the Random Forest Classification method and special polymer-drug descriptors developed previously [25]. Then the *Hybrid strategy* Y-MAX(1)-Y-VAR(1), which proved to be the best on CHEMBL datasets, was applied for AL. The AL cycle was repeated 30 times for every dataset to collect statistics. Since we had only 18 objects in the training set, five of them were selected to form the initial set and with every iteration, one object was added. The performance statistics of the approach was collected on validation set only, i.e., objects left in the modeling set after current set selection. The iterations were carried out until the objects from the pool were exhausted. In Figure 8, a typical plot describing the performance of the selection process is shown (other systems are given in Figure S8-S9 Supporting Materials). It may be noted that median BA of the model built on the *current* set selected by the *Hybrid strategy* was greater than for randomly selected current sets (Figure 8 left). It also provided a good model (mean value BA about 0.83) after only 10 iterations. Moreover, the chance of finding an active polymer at the first iterations of AL procedure was maximal and achieved 50-70%. Notice, that the chance of active polymer selection at random is only 33%. The further gain of chance to find solubilizing polymer can be explained by detection of all active polymers at early stages. Very similar results can be obtained for other datasets (Figures S8 and S9, Supporting Materials). For curcumin (CUR), the hybrid strategy did not show any boost of model performance in comparison with random selection, probably due to high baseline chance to find active compounds (almost 50%). At the same time, one can notice (Figures S8, Supporting Materials) that the chance to find active compound at first iterations of AL cycle (>85%) is much higher than baseline probability.

To summarize, the developed AL strategy can be successfully applied to select polymer compositions that enable high LC for drug molecules. This strategy shows good performance even if only few (5) experimental data is available as initial set.

### 3.3. Validating active learning strategy in an experimental case study

The developed AL strategy and modeling procedure were tested in real-world scenario to find specific polymers that bind specific proteins. It is a complicated case since polymer space is rather big and there is no clear understanding of what features of a polymer can be responsible for protein binding. Also, the experiment is costly and slow, thus the dataset of reasonable size for QSAR modeling can hardly be collected. Here, we searched block copolymers of polyethylene glycol (PEG) and polyamine acids (polylysine - PLKC, polyglutamate - PLE, polyaspartate - PLD) that could bind to the model ASGPR protein at pH 7.4. These polymers are non-toxic and can be applied as scavengers or tracers of proteins or polymer micelles for target delivery.

Initial search space contained 13 different types of polymers (Table 4). Six polymers (4 poly-L-lysine polymers with different numbers of links and different PEG masses, 2 poly-L-glutamate polymers with different numbers of links) were randomly selected, and **K**_**D**_ constants were measured. The values of **K**_**D**_ ranging from 0.51 to 3.36 mM were obtained (Table 4). Polymers with values of **K**_**D**_ less than 1 mM were considered active (i.e., polymers #2 and #12 in Table 4).

**Table 4.** Dataset used to test the AL strategy. K_D_ column contains K_D_ values of initially tested molecules. Six molecules with measured K_D_ values constituted the initial training set. Two molecules (**10** and **11**) were selected after the first iteration of the AL model. The last column K_D_(1^st^ iter) shows the measured K_D_ values for polymers proposed to be tested by the first iteration of the AL strategy

| | 1 block | Mw | 2 block | DP | α-PEG | $K_D$, mM | $K_D(1^{st} \text{ iteration})$, mM |
| --- | --- | --- | --- | --- | --- | --- | --- |
| 1 | PEG | 5k | PLKC | 50 | m | 3.27 |  |
| 2 | PEG | 5k | PLKC | 100 | m | 0.85 |  |
| 3 | PEG | 5k | PLKC | 50 | N3 |  |  |
| 4 | PEG | 5k | PLKC | 30 | m | 1.20 |  |
| 5 | PEG | 5k | PLKC | 10 | m |  |  |
| 6 | PEG | 5k | PLE | 50 | m | 3.36 |  |
| 7 | PEG | 5k | PLE | 50 | N3 |  |  |
| 8 | PEG | 5k | PLE | 100 | m | 1.69 |  |
| 9 | PEG | 5k | PLD | 10 | m |  |  |
| **10** | **PEG** | **1k** | **PLD** | **10** | **m** |  | **1.55** |
| **11** | **PEG** | **1k** | **PLKC** | **100** | **m** |  | **0.02** |
| 12 | PEG | 1k | PLKC | 30 | m | 0.51 |  |
| 13 | PEG | 1k | PLE | 10 | m |  |  |

The α-end of the polyethylene glycol could have a CH<sub>3</sub>O group (m) or an azide group (N<sub>3</sub>).

SiRMS descriptors [52] of the pseudo-small polymer molecules were employed to describe the polymer structure. The *Hybrid active learning strategy* Y-MAX(1)-Y-VAR(1) was applied to develop the classification model: one polymer (#11) with the highest probability to be active and one (#10) with the highest variance were selected at first iteration. According to the surrogate model, polymer #10 was predicted inactive while polymer #11 was predicted active.

The dissociation constant of these polymers was measured experimentally. These predictions were fully confirmed: the predicted active polymer did indeed have a low dissociation constant, whereas polymer #10 was not active. Notice, that polymer #11 had dissociation constant lower (0.023) than the most active candidate (K_D_=0.51), so even the first iteration was successful.

A second AL iteration was undertaken to continue searching for the best performing polymer from the available ones. However, AL system did not find any additional putative active candidates in the set and the calculations were terminated.

To conclude, we observe that the developed AL approach showed its applicability and accuracy both in “virtual” experiments for finding biologically active molecules and polymers with good solubility and in real settings of searching for polymers with a desired ability to bind to a specific protein.

## Conclusion

In this study, we introduced a methodology using classical machine learning methods (classification and regression) in combination with an AL system to discover compounds with the desired properties in a minimum number of the experimental measurements. We benchmarked the performance of the three AL strategies (Y-MAX, Y-VAR, *Hybrid* Y-MAX-Y-VAR) in comparison with *Random selection*, and *Hypothetical perfect strategy* that we proposed for both continuous property and binary datasets. Both regression and classification scenario for AL was explored. In continuous value modeling (regression) AL required selection of quite large number of objects for making informative decisions better than random selection. In case of classification model some AL approaches were found that works well even for small datasets. So, classification modeling is more efficient in AL scenario when only small chemical space exploration is possible. For classification modeling, adding the top candidates (by Y-MAX) has led to the bias in the selected experiments and thereby did not improve the quality of the models over random selection. Adding candidates based on Y-VAR (objects with highest uncertainty are selected at every iteration) causes constant improvement of the model performance, however, the chance to find highly active compound by the respective model is lower than by a model built on random dataset of the same size. Thus, *Hybrid* Y-MAX-Y-VAR strategy was proposed that selects almost half of objects by Y-MAX, and the other half by Y-VAR strategy. *Hybrid* Y-MAX-Y-VAR strategies had benefits of both parent approaches and were capable of continually improving both the performance of classification models and the chance to find highly active compound. The influence of AL method, the size of the initial training set, the way to select the initial data on the performance of AL was thoroughly analyzed in “virtual” experiments. Here, we used datasets of chemical compounds with known biological activities to test AL selection strategies. We found that the formation of the initial dataset from the most diverse objects did not have an advantage over random selection of initial set. For Y-MAX, the model performance obtained using a larger initial training set was higher than those obtained on a smaller initial training set. However, if Y-VAR strategy is applied one can start from smaller dataset and constantly add objects without loss of model accuracy. Also, we found that regression models built even using Y-VAR strategy improve their accuracy much slower with adding AL cycles, than classification models. The former models have lower performance metrics (R^2^) than models built on random datasets. Also, even the best in model performance improvement AL approach, Y-VAR, is quite far from hypothetically perfect AL strategy meaning that there is still a room for improvement and best strategy selection.

The developed AL algorithm was tested on practical cases when AL is really important to use: looking for polymers selectively binding to protein targets. The problem is due to lack of mechanistic insight on binding of polymers to drugs and expensive and slow experimental data collection. Special descriptors reflecting drug and polymer structure were applied for building the surrogate model. The developed AL algorithm has shown its applicability and accuracy both in “virtual” experiments for polymers solubilizing small molecule drugs and in real experiments of searching for polymers with a given ability to bind to a protein.

Previously, AL was primarily used to find quite large datasets for building better model [53] or enhance speed of computational drug design campaigns [54]. Mostly these studies used quite large numbers of data for model building. Here, we show how AL can be efficiently used in case of extremely small datasets, when cost of experiment is high and training sets of reasonable size cannot be collected. It reinforces that tighter interface between experiment and computation opens a door to an extremely time- and resource-efficient design campaigns.

## Acknowledgements

This work was supported by the Russian Science Foundation grant 20-63-46029.

## Abbreviations

AL: active learning
ASGPR: asialoglycoprotein receptor
BA: balanced accuracy
HCC: hepatocellular carcinoma
LC: loading capacity
PLKC: polylysine
PLE: polyglutamate
PLD: polyaspartate
PEG: polyethylene glycol
CUR: curcumin
PTX: paclitaxel
EFV: efavirenz
DEX: dexamethasone
T2A: tanshinone IIA
RFR: Random Forest Regression
RFClf: Random Forest Classification
GPR: Gaussian Process Regression
QRPR: quantitative structure-property relationships
QSAR: quantitative structure-activity relationships
RBF: radial basis function
EF: Enrichment Faction
TPR: true positive rate.

## Supporting Materials

**Fig. S 1.**
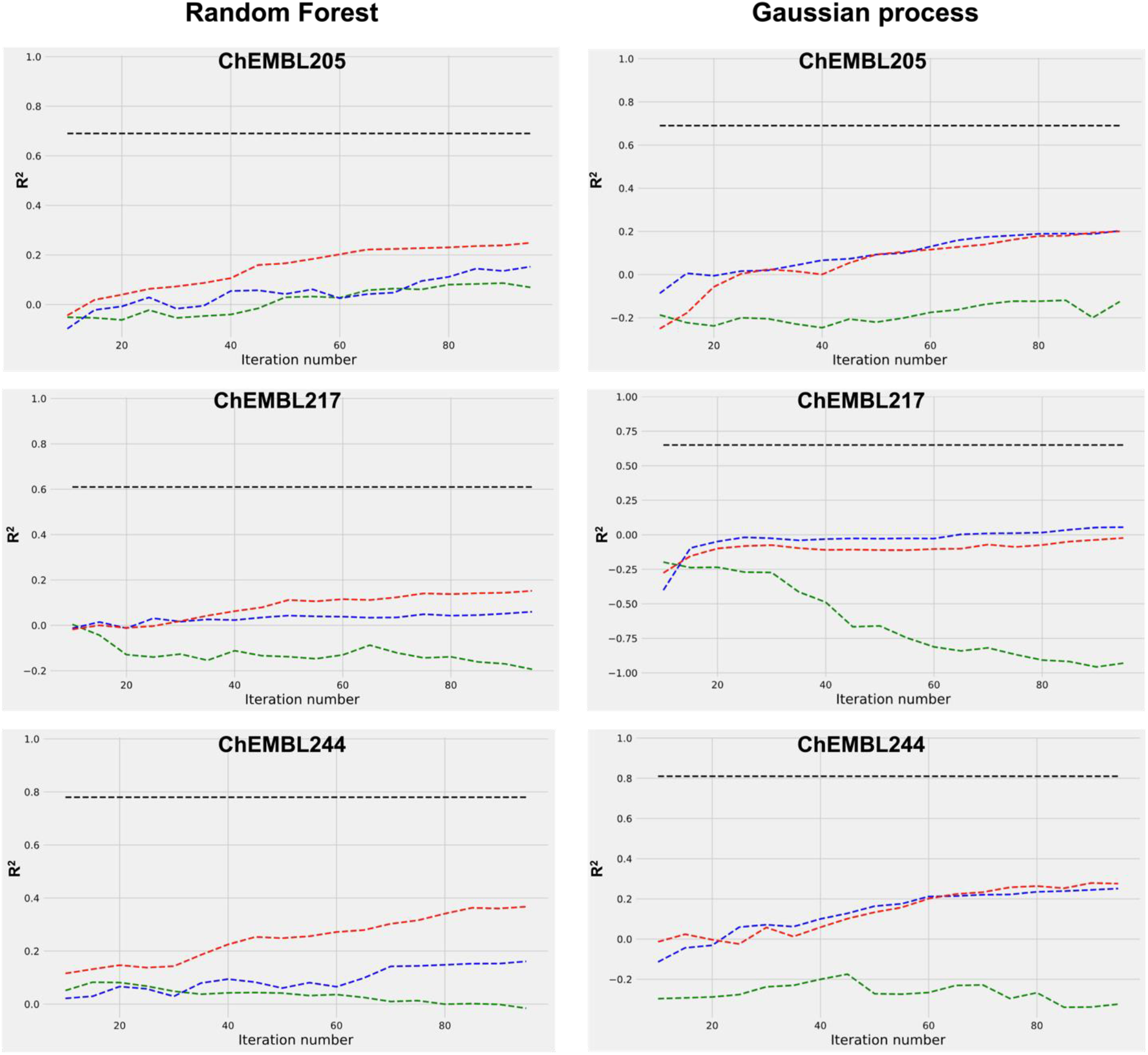
Influence on the quality (R^2^) of RFR and GPR models by different strategies of the AL algorithms. The initial set contains 10 points, 5 objects were added at every AL cycle. Lines correspond to median values over 10 repetitions. Active learning strategies are Y-MAX (green line), Y-VAR (blue line), and random (red line), black dotted line represents performance of the model trained on the entire training data set. Lines correspond to median values over 10 repetitions of AL cycle.

**Fig. S 2.**
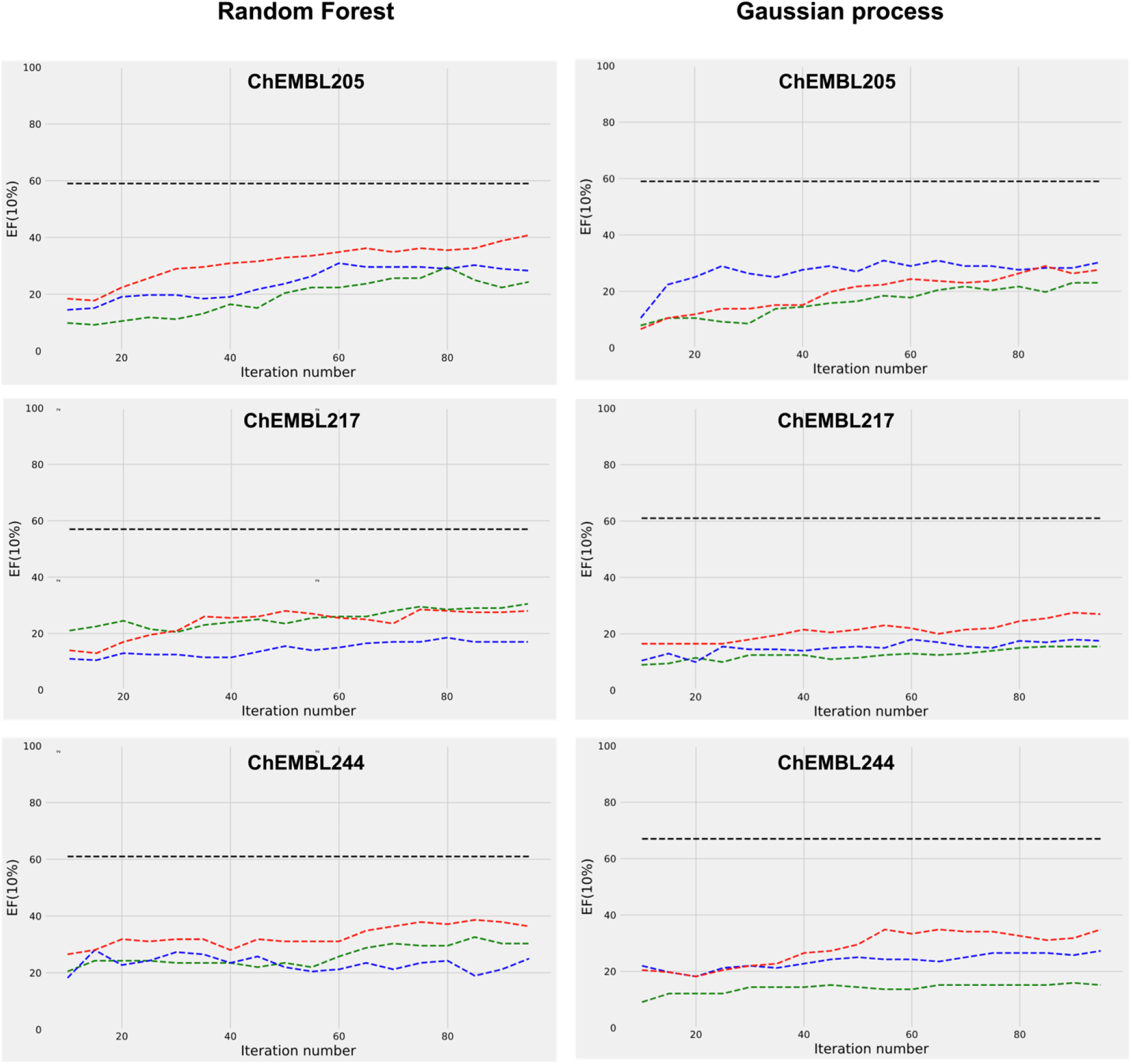
Influence on the quality (EF_10_) of RFR and GPR models by different strategies of the AL algorithms. The initial set contains 10 points, 5 objects were added at every AL cycle. Lines correspond to median values over 10 repetitions. Active learning strategies are Y-MAX (green line), Y-VAR (blue line), and random (red line), black dotted line represents performance of the model trained on the entire training data set. Lines correspond to median values over 10 repetitions of AL cycle.

**Fig. S 3.**
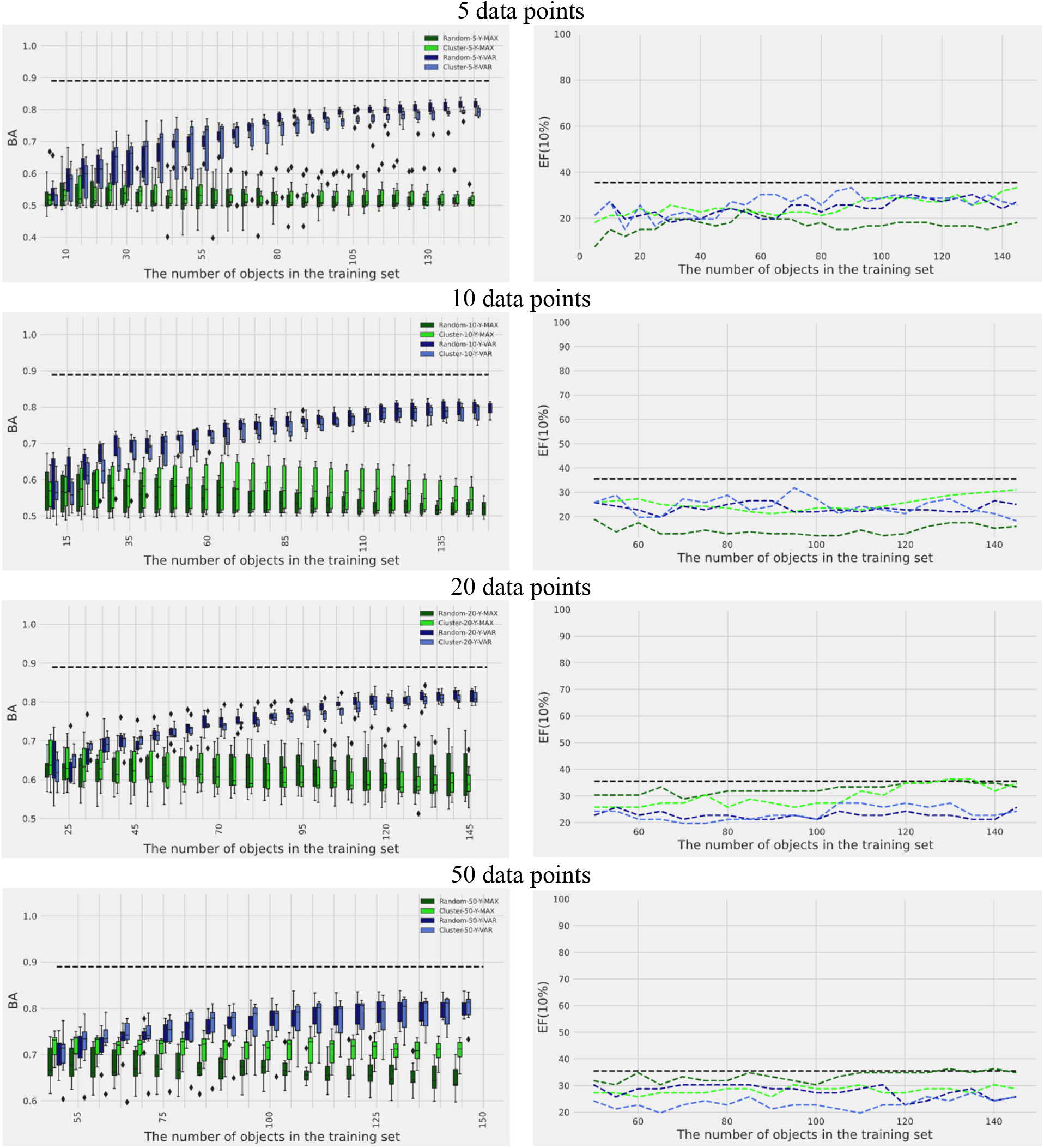
Influence of selection initial dataset strategy. The initial CHEMBL244 dataset consists of 5, 10, 20 and 50 points

**Fig. S 4.**
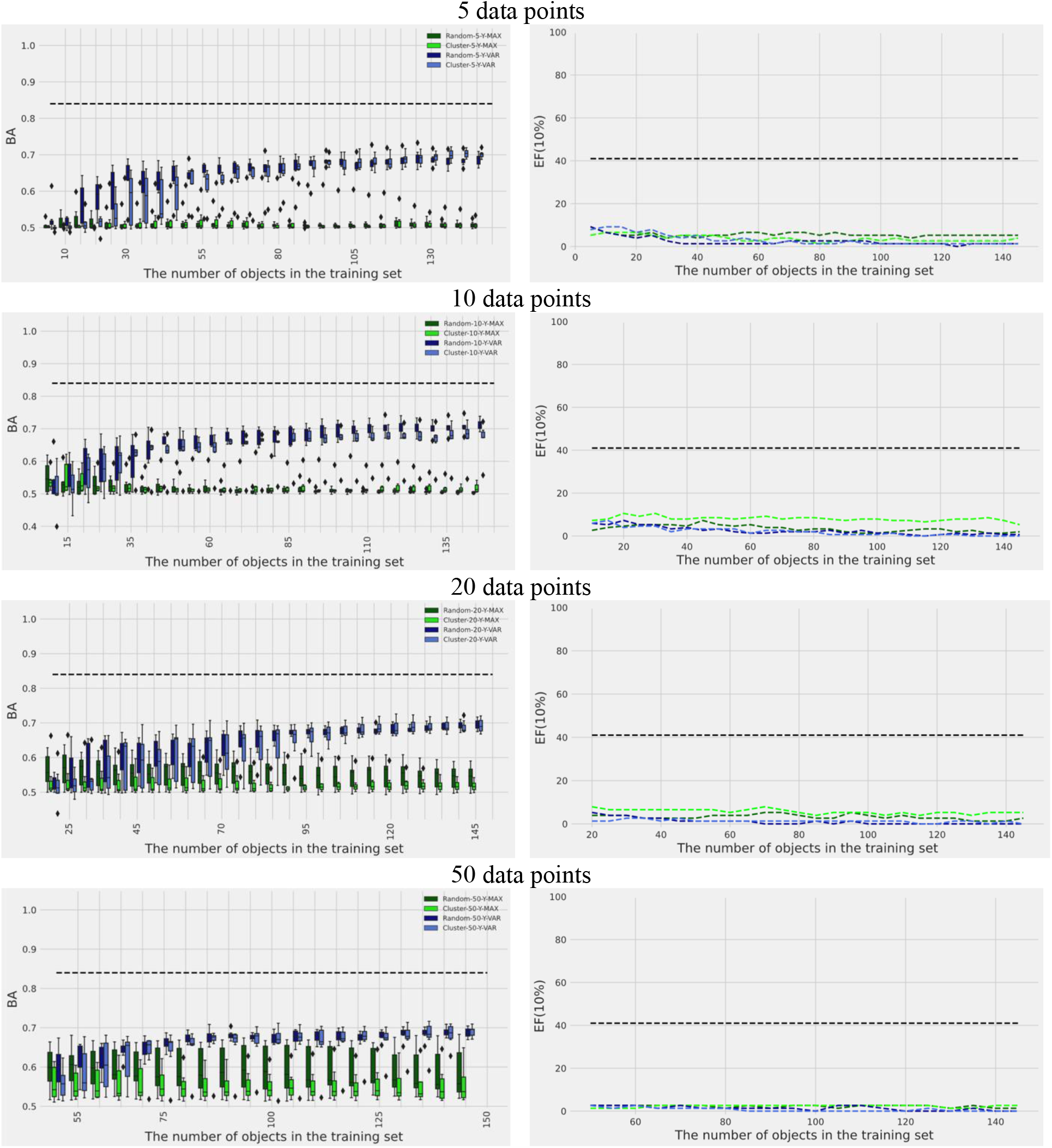
Influence of selection initial dataset strategy. The initial CHEMBL205 dataset consists of 5, 10, 20 and 50 points

**Fig. S 5.**
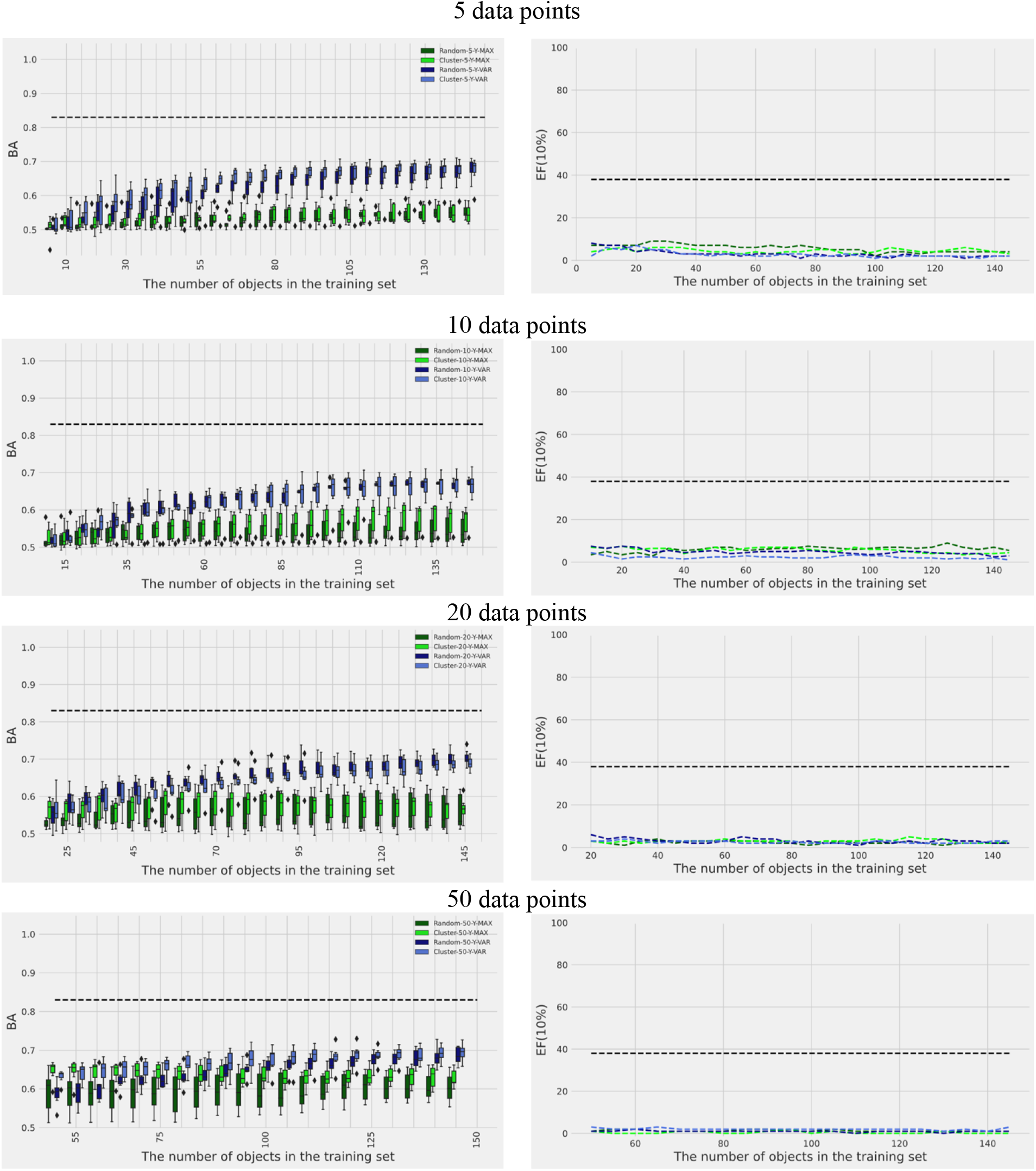
Influence of selection initial dataset strategy. The initial CHEMBL217 dataset consists of 5, 10, 20 and 50 points

**Fig. S 6.**
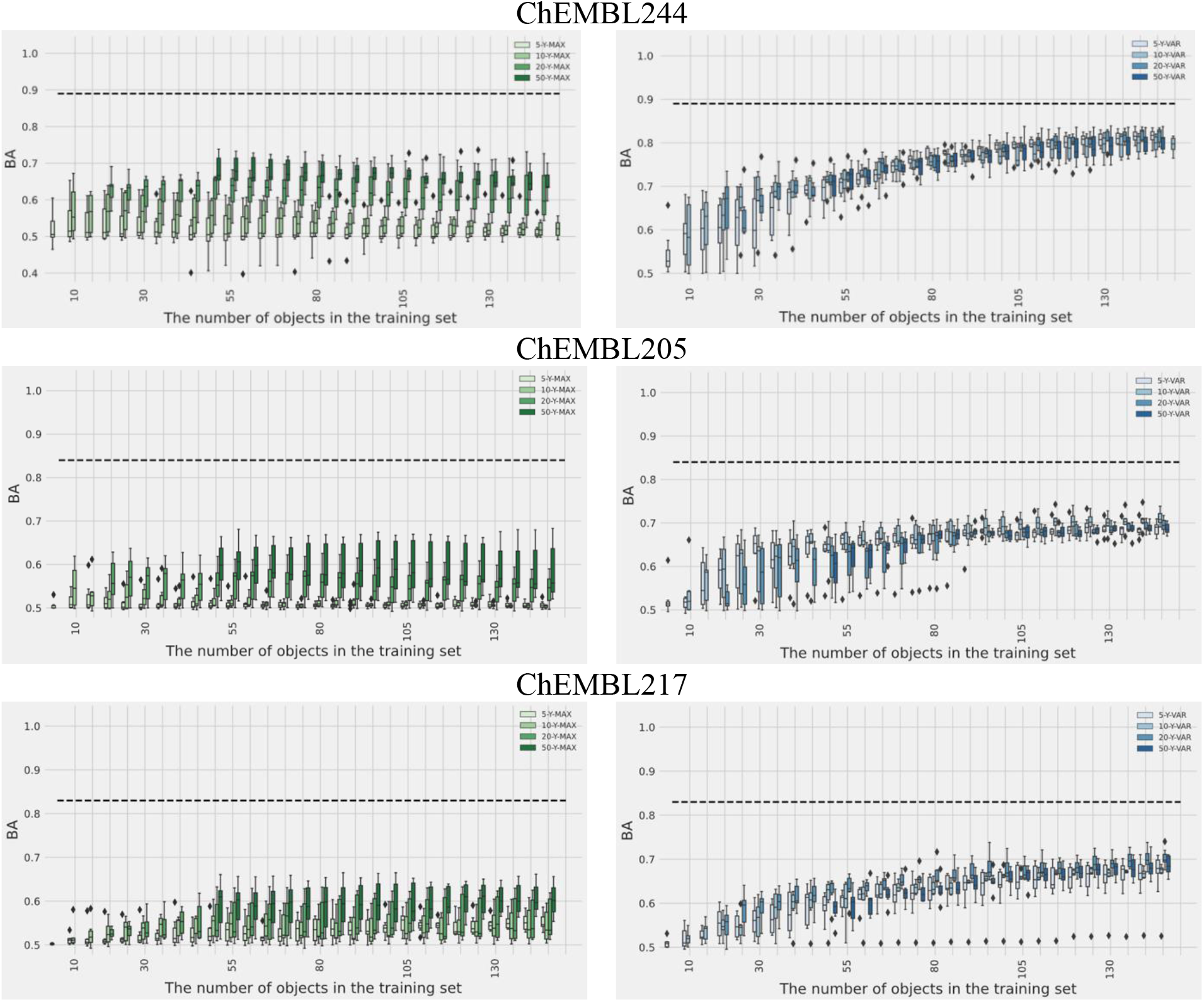
Influence of initial dataset size

**Fig. S 7.**
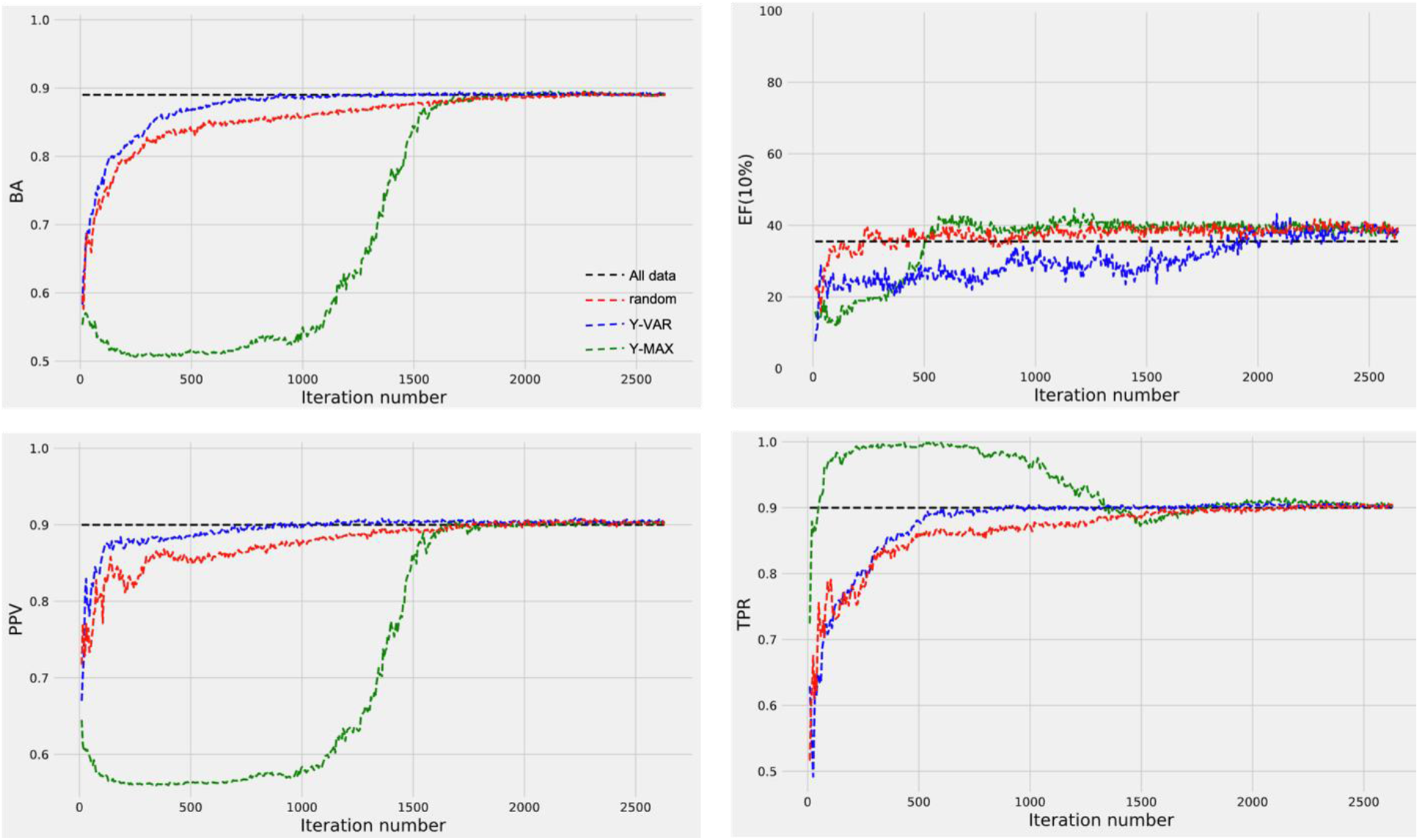
Results of active learning strategies applied to the ChEMBL244 dataset. The initial datasets consist of 10 points. Training the model on all data (black dotted line)

**Table S 1.** The number of active (N «1»)/inactive (N «0») compounds for load capacity (LC) datasets of 5 hydrophobic drugs (Curcumin (CUR), Paclitaxel (PTX), antiretroviral efavirenz (EFV), Dexamethasone (DEX), anti-oxidative tanshinone IIA (T2A)) with amphiphilic polymers investigated under different time. The load capacity values were measured at different time intervals: 0, 5, 10, and 30 minutes after adding the drug to the polymer solution

| Time,<br>min | 0 |  | 5 |  | 10 |  | 30 |  |
| --- | --- | --- | --- | --- | --- | --- | --- | --- |
| Drug | N («0») | N («1») | N («0») | N («1») | N («0») | N («1») | N («0») | N («1») |
| EFV | 14 | 4 | 15 | 3 | 16 | 2 | 15 | 3 |
| CUR | 5 | 13 | 7 | 11 | 7 | 11 | 8 | 10 |
| DEX | 18 | 0 | 0 | 0 | 0 | 0 | 0 | 0 |
| T2A | 18 | 0 | 0 | 0 | 0 | 0 | 0 | 0 |
| PTX | 12 | 6 | 12 | 6 | 13 | 5 | 15 | 3 |

**Table S 2.** Performance of the models on leave-one-out validation for datasets of 3 hydrophobic drugs (Curcumin (CUR), Paclitaxel (PTX), antiretroviral efavirenz (EFV) with amphiphilic polymers investigated under different time. The load capacity values were measured at different time intervals: 0, 5, 10, and 30 minutes after adding the drug to the polymer solution

|  |  | TN | FP | FN | TP | Acc | TPR | TNR | PPV | NPV | BA |
| --- | --- | --- | --- | --- | --- | --- | --- | --- | --- | --- | --- |
| EFV | 0 | 13 | 1 | 3 | 1 | 0.778 | 0.25 | 0.929 | 0.5 | 0.813 | 0.589 |
|  | 5 | 15 | 0 | 3 | 0 | 0.833 | 0 | 1 | - | 0.833 | 0.5 |
|  | 10 | 16 | 0 | 2 | 0 | 0.889 | 0 | 1 | - | 0.889 | 0.5 |
|  | 30 | 13 | 2 | 3 | 0 | 0.722 | 0 | 0.867 | 0 | 0.813 | 0.433 |
| CUR | 0 | 0 | 5 | 2 | 11 | 0.611 | 0.846 | 0 | 0.688 | 0 | 0.423 |
|  | 5 | 4 | 3 | 1 | 10 | 0.778 | 0.909 | 0.571 | 0.769 | 0.8 | 0.740 |
|  | 10 | 3 | 4 | 4 | 7 | 0.556 | 0.636 | 0.429 | 0.636 | 0.429 | 0.533 |
|  | 30 | 5 | 3 | 2 | 8 | 0.722 | 0.8 | 0.625 | 0.727 | 0.714 | 0.713 |
| PTX | 0 | 10 | 2 | 2 | 4 | 0.778 | 0.667 | 0.833 | 0.667 | 0.833 | 0.750 |
|  | 5 | 12 | 0 | 2 | 4 | 0.889 | 0.667 | 1 | 1 | 0.857 | 0.833 |
|  | 10 | 11 | 2 | 4 | 1 | 0.667 | 0.2 | 0.846 | 0.333 | 0.733 | 0.523 |
|  | 30 | 14 | 1 | 3 | 0 | 0.778 | 0 | 0.933 | 0 | 0.824 | 0.467 |

**Fig. S 8.**
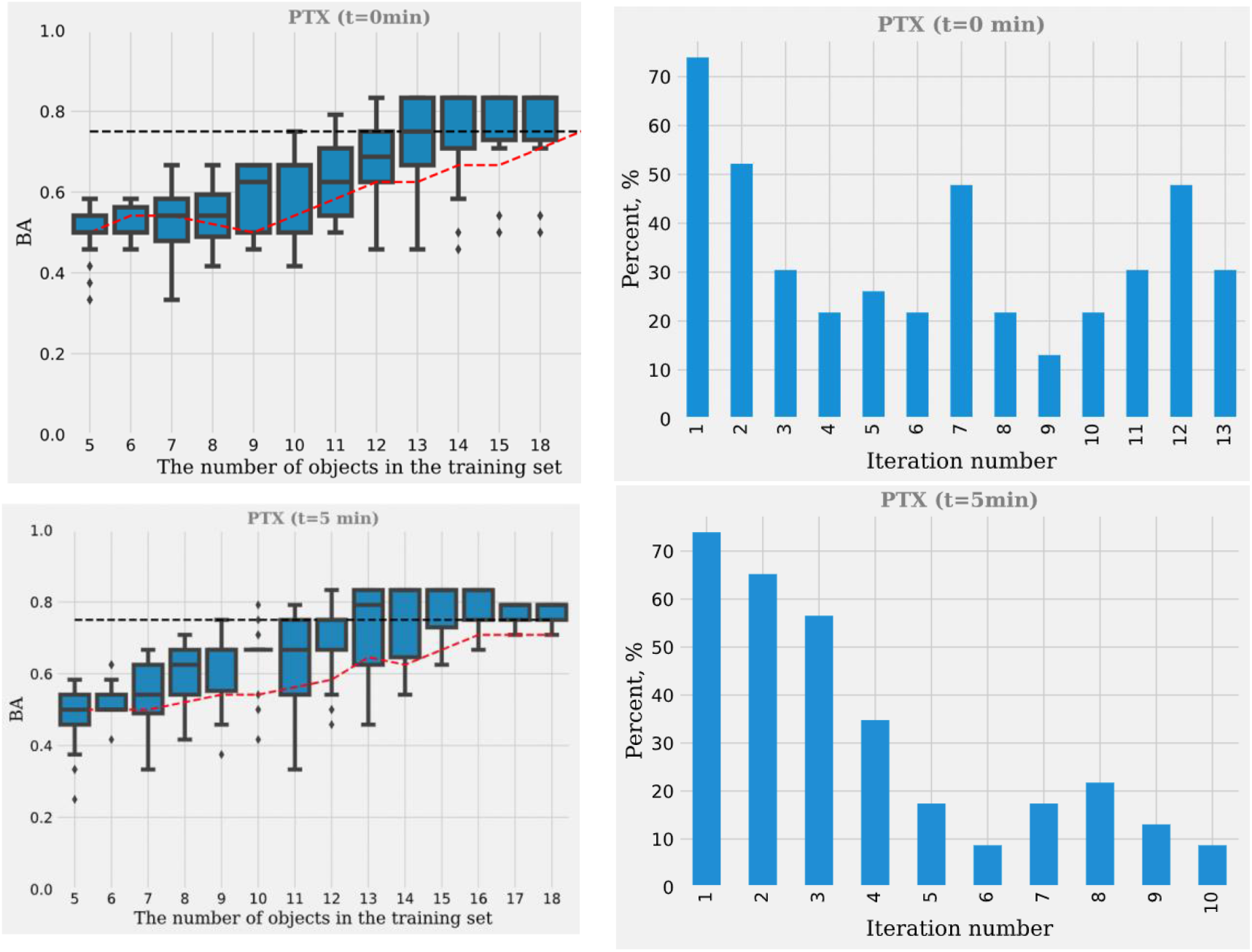
Selection of the optimal polymer for PTX solubilization. Solubilizing ability was studied after 0 and 5 minutes of mixing. On the left - the predictive performance of the model, the BA value is given. On the right - the proportion of active polymer depending on the iterations

**Fig. S 9.**
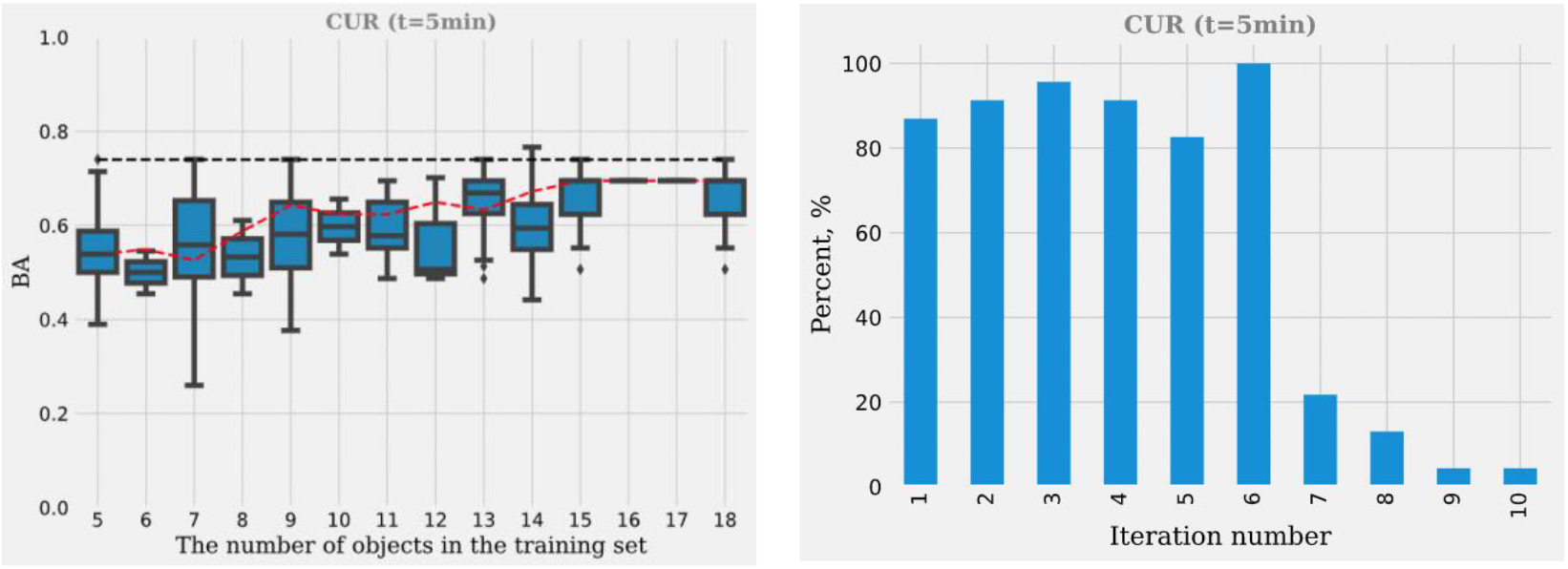
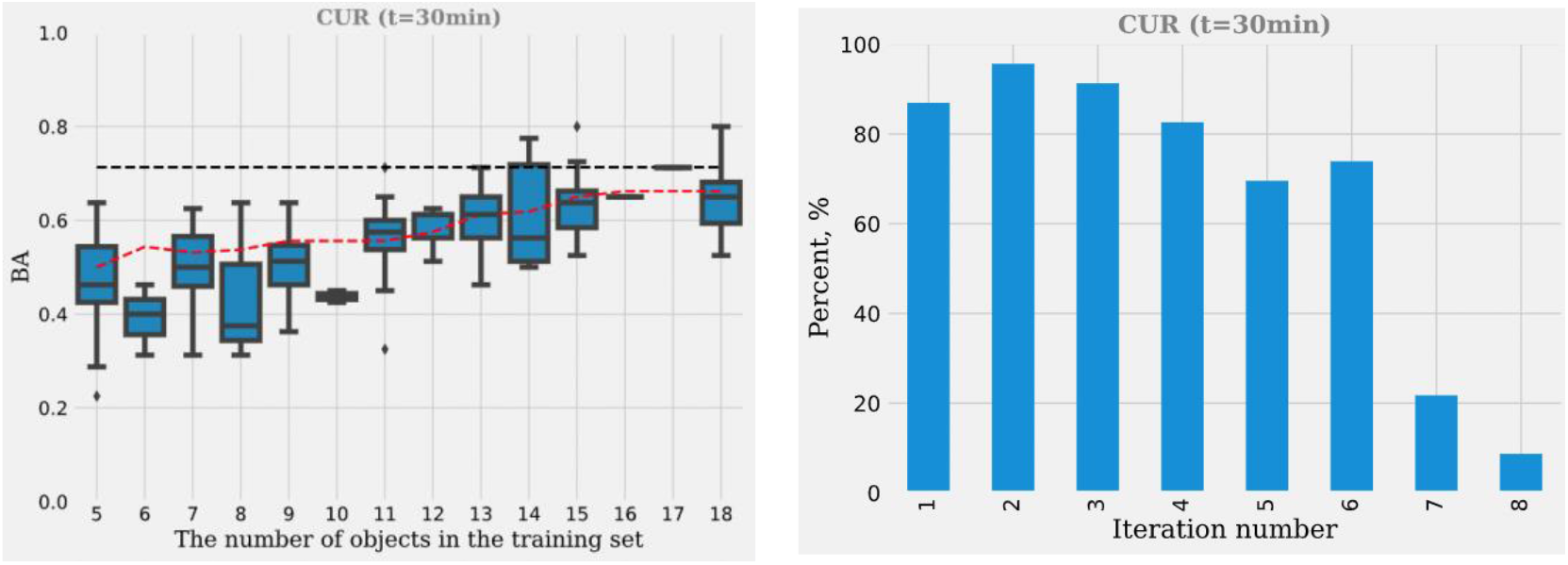
Selection of the optimal polymer for CUR solubilization. Solubilizing ability was studied after 5 and 30 minutes of mixing. On the left - the predictive performance of the model, the BA value is given. On the right - the proportion of active polymer depending on the iterations

## References

1. Baskin I.I. et al. Artificial intelligence in synthetic chemistry: achievements and prospects // Russian Chemical Reviews. 2017. Vol. 86, № 11. P. 1127–1156.

2. Artrith N. et al. Best practices in machine learning for chemistry // Nature Chemistry. 2021. Vol. 13, № 6.

3. Coley C.W., Eyke N.S., Jensen K.F. Autonomous Discovery in the Chemical Sciences Part I: Progress // Angewandte Chemie International Edition. 2020. Vol. 59, № 51.

4. Cherkasov A. et al. QSAR Modeling: Where Have You Beenã Where Are You Going Toã // Journal of Medicinal Chemistry. 2014. Vol. 57, № 12.

5. Muratov E.N. et al. QSAR without borders // Chemical Society Reviews. 2020. Vol. 49, № 11.

6. Settles B. Active Learning // Synthesis Lectures on Artificial Intelligence and Machine Learning. 2012. Vol. 6, № 1.

7. Reker D., Schneider G. Active-learning strategies in computer-assisted drug discovery // Drug Discovery Today. 2015. Vol. 20, № 4.

8. Eyke N.S., Green W.H., Jensen K.F. Iterative experimental design based on active machine learning reduces the experimental burden associated with reaction screening // Reaction Chemistry & Engineering. 2020. Vol. 5, № 10.

9. Kim C. et al. Active-learning and materials design: the example of high glass transition temperature polymers // MRS Communications. 2019. Vol. 9, № 3.

10. Jastrzębski S. et al. Emulating Docking Results Using a Deep Neural Network: A New Perspective for Virtual Screening // Journal of Chemical Information and Modeling. 2020. Vol. 60, № 9.

11. Graff D.E., Shakhnovich E.I., Coley C.W. Accelerating high-throughput virtual screening through molecular pool-based active learning // Chemical Science. 2021.

12. del Rosario Z. et al. Assessing the frontier: Active learning, model accuracy, and multi-objective candidate discovery and optimization // The Journal of Chemical Physics. 2020. Vol. 153, № 2.

13. Kunkel C. et al. Active discovery of organic semiconductors // Nature Communications. 2021. Vol. 12, № 1.

14. Lookman T. et al. Active learning in materials science with emphasis on adaptive sampling using uncertainties for targeted design // npj Computational Materials. 2019. Vol. 5, № 1.

15. Reis M. et al. Machine-Learning-Guided Discovery of ^19^ F MRI Agents Enabled by Automated Copolymer Synthesis // Journal of the American Chemical Society. 2021. Vol. 143, № 42.

16. Smith J.S. et al. Less is more: Sampling chemical space with active learning // The Journal of Chemical Physics. 2018. Vol. 148, № 24.

17. Gubaev K., Podryabinkin E. v., Shapeev A. v. Machine learning of molecular properties: Locality and active learning // The Journal of Chemical Physics. 2018. Vol. 148, № 24.

18. Melnikov A.A. et al. Active learning machine learns to create new quantum experiments // Proceedings of the National Academy of Sciences. 2018. Vol. 115, № 6.

19. Loeffler T.D. et al. Active Learning the Potential Energy Landscape for Water Clusters from Sparse Training Data // The Journal of Physical Chemistry C. 2020. Vol. 124, № 8.

20. Kangas J.D., Naik A.W., Murphy R.F. Efficient discovery of responses of proteins to compounds using active learning // BMC Bioinformatics. 2014. Vol. 15, № 1.

21. Reker D. Practical considerations for active machine learning in drug discovery // Drug Discovery Today: Technologies. 2019. Vol. 32–33.

22. Liu Z.-W., Han B.-H. Evaluation of an Imidazolium-Based Porous Organic Polymer as Radioactive Waste Scavenger // Environmental Science & Technology. 2020. Vol. 54, № 1.

23. Samanta P. et al. Chemically stable microporous hyper-cross-linked polymer (HCP): an efficient selective cationic dye scavenger from an aqueous medium // Materials Chemistry Frontiers. 2017. Vol. 1, № 7.

24. Batrakova E. v. et al. Polymer Micelles as Drug Carriers // Nanoparticulates as Drug Carriers. PUBLISHED BY IMPERIAL COLLEGE PRESS AND DISTRIBUTED BY WORLD SCIENTIFIC PUBLISHING CO., 2006.

25. Alves V.M. et al. Cheminformatics-driven discovery of polymeric micelle formulations for poorly soluble drugs // Science Advances. 2019. Vol. 5, № 6.

26. Harvey H.A. et al. Receptor-mediated endocytosis of Neisseria gonorrhoeae into primary human urethral epithelial cells: the role of the asialoglycoprotein receptor // Molecular Microbiology. 2008. Vol. 42, № 3.

27. Harvey H.A. et al. Gonococcal lipooligosaccharide is a ligand for the asialoglycoprotein receptor on human sperm // Molecular Microbiology. 2000. Vol. 36, № 5.

28. Shi B., Abrams M., Sepp-Lorenzino L. Expression of Asialoglycoprotein Receptor 1 in Human Hepatocellular Carcinoma // Journal of Histochemistry & Cytochemistry. 2013. Vol. 61, № 12.

29. Kanazawa N. Dendritic cell immunoreceptors: C-type lectin receptors for pattern-recognition and signaling on antigen-presenting cells // Journal of Dermatological Science. 2007. Vol. 45, № 2.

30. Rigopoulou E.I. et al. Asialoglycoprotein receptor (ASGPR) as target autoantigen in liver autoimmunity: Lost and found // Autoimmunity Reviews. 2012. Vol. 12, № 2.

31. Becker S., Spiess M., Klenk H.-D. The asialoglycoprotein receptor is a potential liver-specific receptor for Marburg virus // Journal of General Virology. 1995. Vol. 76, № 2.

32. Dotzauer A. et al. Hepatitis A Virus-Specific Immunoglobulin A Mediates Infection of Hepatocytes with Hepatitis A Virus via the Asialoglycoprotein Receptor // Journal of Virology. 2000. Vol. 74, № 23.

33. Mohr A.M. et al. Enhanced colorectal cancer metastases in the alcohol-injured liver // Clinical & Experimental Metastasis. 2017. Vol. 34, № 2.

34. Ueno S. et al. Asialoglycoprotein Receptor Promotes Cancer Metastasis by Activating the EGFR–ERK Pathway // Cancer Research. 2011. Vol. 71, № 20.

35. Pranatharthiharan S. et al. Asialoglycoprotein receptor targeted delivery of doxorubicin nanoparticles for hepatocellular carcinoma // Drug Delivery. 2017. Vol. 24, № 1.

36. Oh H. et al. Galactosylated Liposomes for Targeted Co-Delivery of Doxorubicin/Vimentin siRNA to Hepatocellular Carcinoma // Nanomaterials. 2016. Vol. 6, № 8.

37. Zheng G. et al. Co-delivery of sorafenib and siVEGF based on mesoporous silica nanoparticles for ASGPR mediated targeted HCC therapy // European Journal of Pharmaceutical Sciences. 2018. Vol. 111.

38. Bhingardeve P. et al. Receptor-Specific Delivery of Peptide Nucleic Acids Conjugated to Three Sequentially Linked N -Acetyl Galactosamine Moieties into Hepatocytes // The Journal of Organic Chemistry. 2020. Vol. 85, № 14.

39. Monestier M. et al. ASGPR-Mediated Uptake of Multivalent Glycoconjugates for Drug Delivery in Hepatocytes // ChemBioChem. 2016. Vol. 17, № 7.

40. Thakor D.K., Teng Y.D., Tabata Y. Neuronal gene delivery by negatively charged pullulan– spermine/DNA anioplexes // Biomaterials. 2009. Vol. 30, № 9.

41. Scott L.J. Givosiran: First Approval // Drugs. 2020. Vol. 80, № 3.

42. Fiume L. et al. Liver targeting of antiviral nucleoside analogues through the asialoglycoprotein receptor // Journal of Viral Hepatitis. 1997. Vol. 4, № 6.

43. Plourde R., Wu G.Y. Targeted therapy for viral hepatitis // Advanced Drug Delivery Reviews. 1995. Vol. 17, № 3.

44. Zhang Y. et al. Targeted delivery of atorvastatin via asialoglycoprotein receptor (ASGPR) // Bioorganic & Medicinal Chemistry. 2019. Vol. 27, № 11.

45. Sirtori C.R. The pharmacology of statins // Pharmacological Research. 2014. Vol. 88.

46. Lübtow M.M. et al. Like Dissolves Likeã A Comprehensive Evaluation of Partial Solubility Parameters to Predict Polymer–Drug Compatibility in Ultrahigh Drug-Loaded Polymer Micelles // Biomacromolecules. 2019. Vol. 20, № 8. P. 3041–3056.

47. Zankov D. v. et al. QSAR Modeling Based on Conformation Ensembles Using a Multi-Instance Learning Approach // Journal of Chemical Information and Modeling. 2021. Vol. 61, № 10. P. 4913–4923.

48. RDKit: Open-Source Cheminformatics. http://www.rdkit.org.

49. Pedregosa F. et al. Scikit-learn: Machine Learning in Python. 2012.

50. Scikit-Learn User Guide. Available online: https://scikit-learn.org/stable/_downloads/scikit-learn-docs.pdf.

51. Rasmussen, C.E.; Williams, C.K.I. Gaussian Processes for Machine Learning; MIT Press: Cambridge, MA, USA, 2006; ISBN 026218253X. Available online: http://www.gaussianprocess.org/gpml/chapters/RW.pdf.

52. Muratov E.N. et al. Per aspera ad astra : application of Simplex QSAR approach in antiviral research // Future Medicinal Chemistry. 2010. Vol. 2, № 7. P. 1205–1226.

53. Smith J.S. et al. Less is more: Sampling chemical space with active learning // The Journal of Chemical Physics. 2018. Vol. 148, № 24.

54. Yang Y. et al. Efficient Exploration of Chemical Space with Docking and Deep Learning // Journal of Chemical Theory and Computation. 2021.

